# Statistical Characterization of Cortical-Thalamic Dynamics Evoked by Cortical Stimulation in Mice

**DOI:** 10.1101/2025.04.04.647222

**Authors:** Diana Nigrisoli, Simone Russo, Ruggero Freddi, Nicolas Seseri, Stefania Corti, Linda Ottoboni, Riccardo Barbieri

## Abstract

**Objective:** Statistical models are powerful tools for describing biological phenomena such as neuronal spiking activity. Although these models have been widely used to study spontaneous and stimulated neuronal activity, they have not yet been applied to analyze responses to electrical cortical stimulation. In this study, we present an innovative approach to characterize neuronal responses to electrical stimulation in the mouse cortex, providing detailed insights into cortical-thalamic dynamics.

**Approach:** Our method applies Mixture Models to analyze the Peri-Stimulus Time Histogram of each neuron, predicting the probability of spiking at specific latencies following the onset of electrical stimuli. By applying this approach, we investigated neuronal responses to cortical stimulation recorded from the motor cortex, somatosensory cortex, and sensorimotor-related thalamic nuclei in the mouse brain.

**Main results:** The characterization approach achieved high goodness of fit, and the model features were leveraged by applying machine learning methods for stimulus intensity decoding and classification of brain regions to which a neuron belongs given its response to the stimulus. The Random Forest model demonstrated the highest F1 scores, achieving 92.86% for stimulus intensity decoding and 84.35% for brain zone classification.

**Significance:** This study presents a novel statistical framework for characterizing neuronal responses to electrical cortical stimulation, providing quantitative insights into cortical-thalamic dynamics. Our approach achieves high accuracy in stimulus decoding and brain region classification, providing valuable contributions for neuroscience research and neuro-technology applications.

## 1 Introduction

Understanding neural activity is crucial for advancing our knowledge of brain function, developing innovative treatments for neurological disorders, and implementing neural control in brain-computer interfaces (BCIs) [1]. BCIs enable bidirectional communication between the brain and external actuators, with high potential for assisting individuals with disabilities and controlling prosthetic limbs. Neurons transmit and process information inside the brain, therefore understanding neural activity is a crucial step for performing complex tasks with BCIs [2]. Neural activity can be characterized through statistical models, which solve the encoding-decoding problem. The encoding step calculates the probability of a neuron’s activity given a stimulus or a condition, while the decoding problem determines the probability of the stimulus or condition given the neuron’s activity [3]. A common approach for characterizing neuronal activity leverages Generalized Linear Models to estimate firing probability combining point process theory and inhomogeneous Poisson distribution [3, 4, 5, 6, 7]. Capturing spiking activity with higher accuracy requires more complex frameworks, incorporating renewal processes and a transformation function that links spike history, external stimuli, and model parameters [8, 9]. Alternatively, leveraging features and statistics derived from neural data representations, such as the Inter-spike Interval (ISI) histogram [10] and the Peri-Stimulus Time Histogram (PSTH) [11, 12], can offer a quantitative insight on neuronal activity. These approaches have been used extensively to elucidate the properties of both spontaneous neuronal activity and activity evoked by different types of stimuli or pathological perturbations. Examples include auditory stimuli [13, 14], as well as electrical stimulation to auditory nerve fibers [15]. Visual and proprioception stimuli [8, 9] have also been studied. In the context of pathological conditions, research includes analysis of neural activity in neural networks with gene mutation for Parkinson’s disease [10], modeling neuronal activity in Parkinson’s patients [16] and modeling epilepsy [17]. However, to our knowledge, these approaches have not yet been applied to characterize responses to electrical cortical stimulation.

In this study, we introduce a new framework to characterize neuronal responses evoked by the electrical stimulation of the mouse cortex, offering detailed insights into cortical-thalamic dynamics. Indeed, electrical cortical stimulation is applied in research and medical settings in order to investigate brain function and map neural pathways [18] or to diagnose and treat certain neurological disorders [19]. Moreover, electrical stimulation can be used in BCIs to enable bidirectional communication between the brain and external devices (e.g. restoring vision via cortical visual prosthetics [20]). Here we apply our framework to a dataset of neuronal responses evoked by cortical stimulation in mice published by Claar et al. [21]. They analysed this dataset and showed how cortico-thalamo-cortical (CTC) interactions modulate electrically evoked EEG responses in mice [22]. Claar et al. investigated the neural circuits in mice by delivering deep cortical stimulation and recording ipsilateral neural activity through EEG and Neuropixels probes (Figure 1 A). The results show that during wakefulness the cortical stimulation evokes in the cortical layer a short spiking response (early response), followed by an off period and a rebound spiking activity (late response). Therefore, the response can be divided into three temporal windows following the stimulus: 2–25 ms (early response), 25–150 ms (off period), and 150–300 ms (late response). The thalamus exhibits a similar triphasic pattern in which the last phase corresponds to late burst spiking. Furthermore, the late thalamic activity is temporally linked to EEG signals evoked in the same time window, suggesting that cortico-thalamic-cortical (CTC) interactions contribute to the generation of evoked EEG responses [22]. A second study conducted by Russo et al. [23] demonstrated that the delayed aspect of the EP in mice is caused by the thalamic hyperpolarization and rebound (Figure 1 B). The intensity of the late response is linked to the bursting rate and synchronization of thalamic neurons, which are influenced by the individual’s behavioral condition and thalamocortical synaptic connections.

**Figure 1:**
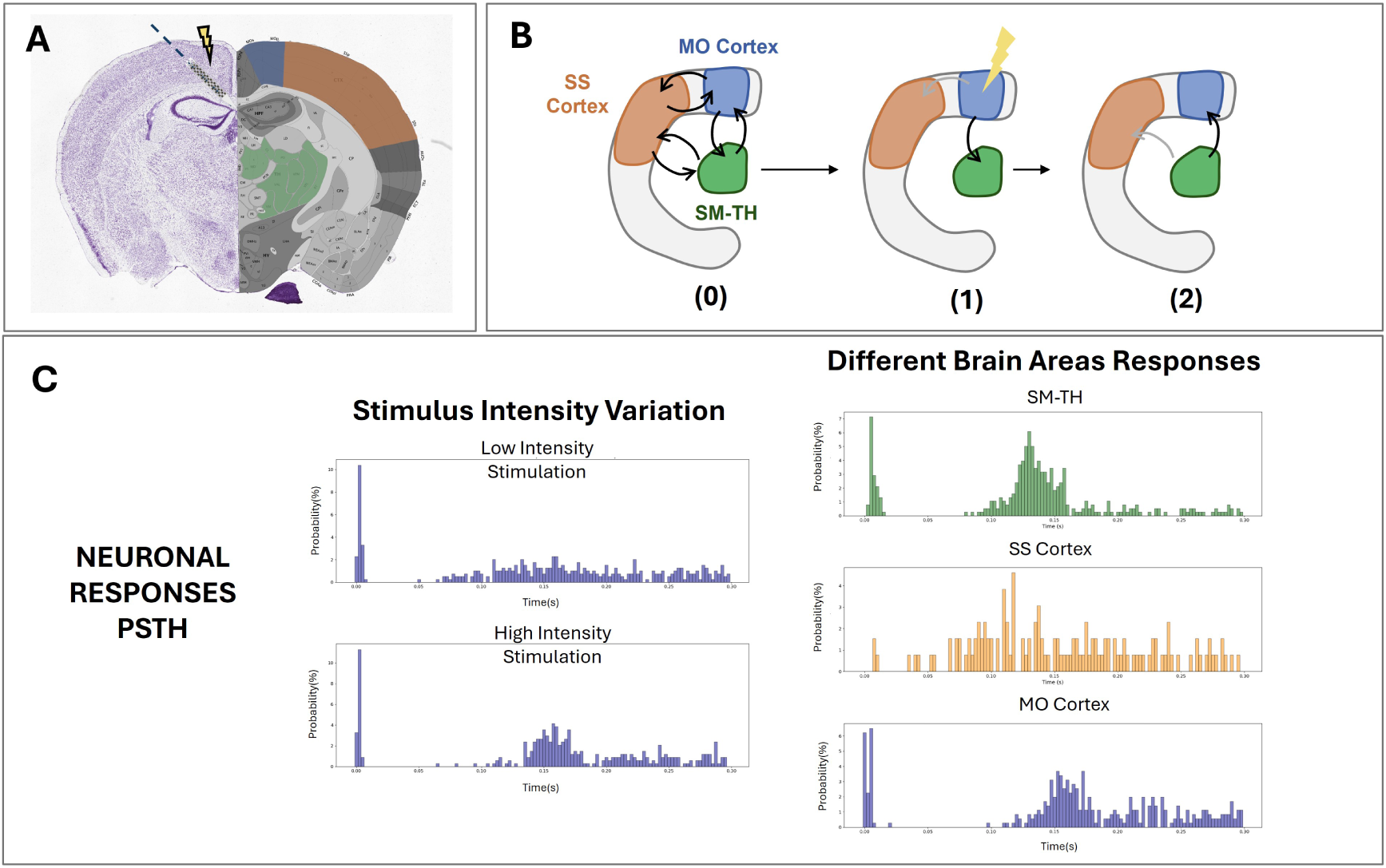
(A) Coronal atlas slice [25] showing the placement of the bipolar stimulating electrode (represented as a lightning) and a Neuropixels probe (blue line), modified from [22] (B) Diagram of thalamocortical loops between motor (MO) cortex, sensorimotor (SS) cortex and sensorimotor related thalamic nuclei (SM-TH) where the MO cortex is electrically stimulated. The interactions between the stimulated cortex and thalamus (0) involve a dynamic cortico-thalamo-cortical (CTC) loop in which cortical stimulation elicits early responses (1) and late responses (2), with the thalamus contributing to the delayed rebound through burst spiking [22, 23]. Grey arrows indicate hypothetical interactions between unstimulated cortex and either stimulated cortex and SM-TH. (C) Neuronal responses PSTH showing the same MO cortex neuron stimulated with low and high current intensity (left) and three representative PSTH evoked in SMTH, SS and MO cortex by the stimulation of MO cortex.

To test our framework, we analyzed the neuronal responses recorded from the motor (MO) cortex, somatosensory (SS) cortex, and sensorimotor-related thalamic (SM-TH) nuclei, from both MO and SS deep stimulation. The temporal dynamics of neuronal responses to stimulation can be represented by the PSTH, which is linearly related to the average firing rate of the triggered spike train evaluated for every bin [24]. Our framework consists in implementing

Dirichlet Mixture Models (DMM) to model the PSTH and, therefore, predict the probability of spike occurrences at specific latencies following electrical stimulus onset. In general, when developing a neural characterisation model, the following criteria must be taken into account [3]. The model should fit within the constraints of the available data. Additionally, it should not be too complex and should be interpretable from a physiological perspective. Finally, it should align with existing knowledge about the underlying physiology of the system. This analysis pipeline is designed to provide a detailed description of the response pattern of each neuron and, thus, differentiate between types of neuronal responses across stimulation intensities and brain area by combining flexibility and robustness. Our approach not only characterizes the neuronal response patterns but also facilitates the decoding of stimulus intensity and classification of brain regions based on their response to stimuli.

This study has three primary objectives. First, we devise a robust statistical framework for characterizing neuronal responses to electrical stimulation. Second, we apply this framework to analyze neuronal responses in the cortex and thalamus, allowing for a detailed description of each neuron’s response pattern and distinguishing between different types of neuronal responses. Furthermore, we seek to exploit this model using machine learning techniques to decode the stimulus intensity and classify brain regions to which a neuron belongs given its response to the stimulus. Lastly, we use the results of this framework to provide insights into cortical-thalamic dynamics following cortical electrical stimulation. In particular, we analyze the differences in response patterns to varying stimulation intensities (Figure 1 C). Finally, we investigate the differences between the activity of one cortical area (e.g., SS cortex) while directly stimulating another (e.g., MO cortex) (Figure 1 B,C). By accurately modeling and decoding neuronal activity, we can enhance our comprehension of the underlying mechanisms of brain function, offering significant insights on the patterns engaged by cortical stimulation.

## 2 Materials and Methods

This section details the methodologies and procedures employed in our study to analyze the neural data and evaluate model performance. In particular, we employ the encoding/decoding paradigm, which is central to neural characterization. The encoding step involves calculating the probability of a neuron’s activity in response to a given stimulus or condition, while the decoding step involves determining the probability of the stimulus or condition based on the neuron’s activity described by the encoding model [3]. We begin by describing the stimulation protocols and data acquisition methods, which include the collection and processing of neural activity data as outlined in Claar et al. [22]. We then explain the data selection process, focusing on the criteria used to identify significant neuronal responses and ensure the data’s relevance for subsequent analysis. Next, we introduce the probabilistic model used for analyzing spiking activity, highlighting the use of maximum a posteriori estimation to determine model parameters and optimize fitting accuracy. Finally, we describe the evaluation metrics and methods used to assess both model adherence to observed data and the adequacy of its parameters for decoding. This included training machine learning algorithms to decode stimulus characteristics (intensity and brain area) from spike distribution patterns based solely on the model’s parameters. Each subsection provides a comprehensive overview of the specific aspects of our methodology, ensuring a thorough understanding of the approaches used to achieve our research objectives.

### 2.1 Stimulation Protocol and Data Acquisition

We here summarize the stimulation protocols and data acquisition methods used in the study. Detailed descriptions of probe placement, stimulation parameters, and data processing are provided based on the work of Claar et al. [22]. The data related to the mice neural activity that we consider are also from of Claar et al. [22] and can be downloaded from the DANDI archive [21]. In summary, up to three Neuropixels probes [26] were used for data recording on each subject and were inserted targeting motor (MO), anterior cingulate, somatosensory (SS), visual cortex areas, and the thalamic nuclei. The data was acquired with a sampling frequency of 30 kHz and high-pass filtered at 500Hz. For cortical stimulation a custom bipolar platinum-iridium stereotrode (Microprobes for Life Science, Gaithersburg, MD, USA) made by two parallel monopolar electrodes was used with a vertical offset of 300 µm between the two tips. For each subject, stimuli were released in one of two areas of the cortex: the secondary motor area or in the primary somatosensory area. Up to 120 biphasic, charge-balanced, cathodic-first current pulses (200 *µ*s per phase, 3.5–4.5 s interstimulus interval) were administered at each of three different current intensities (i.e. low, medium, high), totaling 360 pulses. For the purposes of this study, we consider *low* and *high* stimulation intensities. Eight mice received electrical stimulation in the MO cortex and seven mice received it in the SS cortex, while one mouse received stimulation in both areas during two different recording sessions [22]. Spike sorting step was performed by Claar et al. with the Kilosort algorithm [27]. For further details see [22].

### 2.2 Data Selection and Settings

This section provides an overview of the neurons included in the dataset and details the criteria used to select neurons for analysis, focusing on those whose post-stimulus activity significantly differs from baseline.

The dataset comprises both regular spiking (RS) and fast spiking (FS) neurons, identified by their spike waveform duration (RS: *>*0.4 ms; FS: ≤0.4 ms) [28, 29]. Our analysis focuses on RS neurons within the MO cortex, SS cortex, and SM-TH, which include various nuclei such as the anteroventral, central lateral, mediodorsal, posterior, reticular, ventral anterior-lateral, ventral posterolateral, ventral posteromedial, and ventral medial nuclei. Only spike data related to the wakefulness phase was considered.

To ensure robust data selection, we applied specific criteria for response significance and activity representation. Neurons were considered responsive based on the following pipeline:

1. Compute the PeriStimulus Time Histogram (PSTH) for pre-stimulus spikes (psth1) and post-stimulus spikes (psth2) within defined windows (i.e. (-0.3, 0 s) and (0, 0.3 s) with respect to the stimulus onset).
2. Calculate the mean (*m*) and standard deviation (std) of psth1.
3. Set the threshold as threshold = *m*+4×std. This threshold maximizes the identification of neurons that show a clear change in firing relative to baseline activity while simultaneously reducing false positives.
4. Mark a neuron as responsive if at least one bin in psth2 exceeded this threshold.

The bin width was set to 2.5 ms across all PSTHs to balance resolution and activity representation. Additionally, a minimum of 50 spikes across stimulation trials was required to ensure adequate representation of neuronal activity.

This selection processes isolated neurons with activity levels that clearly diverge from baseline, supporting effective analysis by subsequent statistical models.

### 2.3 Model Characterization

To characterize neural dynamics, we use a specific family of point processes (PPs) modeling spike occurrences. The Dirichlet Mixture Model (DMM) is employed to fit the data distribution, using uniform priors for weights to avoid bias. Parameters are then estimated using maximum a posteriori (MAP), and the goodness of fit of the model is assessed using the Kolmogorov-Smirnov (KS) test.

A PP is a binary stochastic process that models irregular events occurring in continuous time or space. In the context of neuronal activity, time is divided into short intervals, during which each neuron either spikes (1) or remains silent (0). This framework captures the probabilistic variation in spike occurrences over time, and modeling the precise timing of spikes requires estimating the associated probability functions [24].

We denote *f* (*t*) as the probability of observing a spike at time *t*, with time *t* anchored to the stimulus onset. To model the distribution of spike timings, we use a DMM defined as:

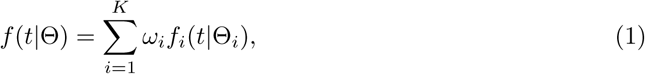

where *f_i_*(*t*|Θ*_i_*) are the component distributions, Θ*_i_* are their parameters, and *ω_i_* are the weights in the convex combination. The model parameters are summarized as Θ = (*ω*_1_*, …, ω_K_,* Θ_1_*, …,* Θ*_K_*).

The weights *ω* are constrained by:

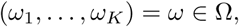

where

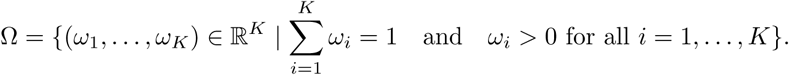

We model the distribution of weights using the Dirichlet distribution:

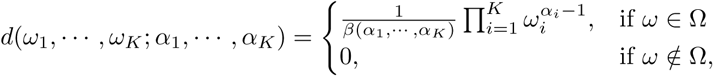

where *α* = (*α*_1_*, …, α_K_*) are the concentration parameters, and *β*(*α*) is the normalizing constant:

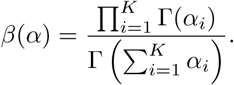

The Dirichlet distribution provides flexibility in modeling the weight distribution according to prior beliefs [30].

For parameter estimation, we use MAP estimation to find the parameters that maximize the posterior distribution:

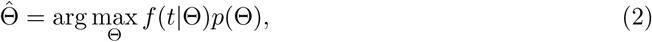

where *p*(Θ) is the prior distribution over Θ, and *f* (*t*|Θ) is the likelihood.

Uniform distributions are used for both the prior *p*(Θ) and the weights *ω_i_* to avoid bias. Specifically, setting *α_i_* = 1 for all *i* results in a uniform distribution over Ω, ensuring that no preference is given to any particular set of weights and allowing the data to drive the estimation process.

### 2.4 Encoding: Models definition and Prior Probabilities

Here, we outline the DMM selection, detailing the tested Gaussian and inverse Gaussian combinations used to approximate both early and late phases of neural responses in the PSTH data.

Given the bi-phasic nature of the neural response—comprising an early phase (0-0.05 s) and a late phase (0.05-0.3 s)—and based on visual inspection of the PSTHs, it was determined that combinations of two or three Gaussian and inverse Gaussian distributions would be appropriate for approximating *f* (*t*) and, therefore, the PSTH. Specifically, the early response was modeled using a Gaussian distribution, while the late response was approximated using various combinations of Gaussian and inverse Gaussian distributions (Table 1). The prior probabilities *p*(Θ) of the parameters of these curves were defined by uniform distributions (∼ *U* (min, max)). The extremes min, max were set so as to temporally constrain the mean *µ* of the curves to certain intervals but leaving wide room for *σ* and *λ* in order to shape the curves correctly (Table 2).

**Table 1:** Summary of the Tested Models for Mice Neurons. G stands for Gaussian and InvG stands for Inverse Gaussian.

| Model | Early Response | Late Response |
| --- | --- | --- |
| 2 G | 1 G | 1 G |
| 1 G 1 InvG | 1 G | 1 InvG |
| 3 G | 1 G | 2 G |
| 1 G 2 InvG | 1 G | 2 InvG |
| 2 G 1 InvG | 1 G | 1 G and 1 InvG |

**Table 2:** Prior Probabilities for Tested Model Parameters

| Model | Parameters 1st curve | Parameters 2nd curve | Parameters 3rd curve |
| --- | --- | --- | --- |
| 2G | $\mu_1 \sim U(0, 0.05),$<br>$\sigma_1 \sim U(10^{-6}, 1)$ | $\mu_2 \sim U(0.05, 0.3),$<br>$\sigma_2 \sim U(10^{-6}, 1)$ | - |
| 1G 1InvG | $\mu_1 \sim U(0, 0.05),$<br>$\sigma_1 \sim U(10^{-6}, 1)$ | $\mu_2 \sim U(0.05, 0.3),$<br>$\lambda_2 \sim U(10^{-6}, 1)$ | - |
| 3G | $\mu_1 \sim U(0, 0.05),$<br>$\sigma_1 \sim U(10^{-6}, 1)$ | $\mu_2 \sim U(0.05, 0.3),$<br>$\sigma_2 \sim U(10^{-6}, 1)$ | $\mu_3 \sim U(\mu_2, 0.3),$<br>$\sigma_3 \sim U(10^{-6}, 1)$ |
| 1G 2InvG | $\mu_1 \sim U(0, 0.05),$<br>$\sigma_1 \sim U(10^{-6}, 1)$ | $\mu_2 \sim U(0.05, 0.3),$<br>$\lambda_2 \sim U(10^{-6}, 1)$ | $\mu_3 \sim U(\mu_2, 0.3),$<br>$\lambda_3 \sim U(10^{-6}, 1)$ |
| 2G 1InvG | $\mu_1 \sim U(0, 0.05),$<br>$\sigma_1 \sim U(10^{-6}, 1)$ | $\mu_2 \sim U(0.05, 0.3),$<br>$\sigma_2 \sim U(10^{-6}, 1)$ | $\mu_3 \sim U(0.05, 0.3),$<br>$\lambda_3 \sim U(10^{-6}, 1)$ |

The accuracy of each model is measured via the KS test, which computes the distance between the cumulative distribution function (CDF) of the model to that of the observed data [31]. A small KS distance and a p-value near 0.95 indicate a good fit, with the KS plot ideally showing a 45° slope and the model’s CDF remaining within the 95% confidence interval [10].

### 2.5 Decoding Tasks

Here we present decoding tasks that assess the model’s ability to effectively represent and discriminate neural responses.

After selecting an accurate model through the KS test, we validated its efficacy by assessing how well its parameters performed in specific decoding tasks [10, 3]. These tasks were designed to evaluate the model’s ability to capture meaningful features of neuronal responses:

1. **Stimulus Intensity Decoding:** This binary classification task distinguishes between low and high stimulus intensities based on neuronal responses. Methods are detailed in subsection 3.4.
2. **Brain Area Decoding:** This multiclass classification task identifies the neuron’s brain region in response to stimuli applied to either the MO cortex or the SS cortex, with classes for ”MO cortex,” ”SS cortex,” and ”SM-TH.” See subsection 3.5 for details.

These decoding tasks assess the model’s capacity to retain information on stimulus characteristics, with high performance indicating an accurate representation of neuronal activity. Additionally, the model’s interpretability is confirmed if it aligns with physiological knowledge by revealing differences in neuronal responses based on location. An effective model, therefore, will demonstrate both strong decoding accuracy and physiological relevance.

#### 2.5.1 Statistical Evaluation of Model Parameters

This section explains the statistical methods used to determine if the model parameters are suitable for distinguishing between stimulus intensities and classifying brain areas. It focuses on analyzing neural responses to different stimuli and examining their statistical differences.

For intensity decoding, we analyzed neurons from the directly stimulated regions (MO cortex or SS cortex) that responded to both stimulus intensities. The model parameters Θ were compared to determine their discriminative power between stimulus intensities. Before applying ML methods, statistical tests were performed to assess significant differences between data classes. Specifically, Wilcoxon signed-rank tests were applied, with Bonferroni correction employed to account for multiple comparisons.

For brain area classification, neurons were grouped by location (SS cortex, MO cortex, or SM-TH) considering their response to the highest intensity stimulus. Statistical tests were conducted to assess significant differences in feature distributions across these regions. Mann-Whitney tests were utilized, with Bonferroni correction applied to adjust for multiple comparisons.

#### 2.5.2 Machine Learning Models for Classification

The performance of seven supervised ML models—K-Nearest Neighbors, Logistic Regression, Support Vector Machines, Decision Tree, Random Forest, AdaBoost, and XGBoost—was assessed for classifying both stimulus intensity and brain area. These models were trained using features derived from the parameter set Θ, which represents the defining characteristics of the assumed underlying probabilistic distribution for the neural process. Specifically, Θ includes parameters such as the means and standard deviations values associated with Gaussian components within a Dirichlet Mixture Model. As these parameters encapsulate the distribution’s structure, they serve as the core characteristics required to identify the process and are expected to be sufficient for reliable inference.

Hyperparameter optimization was performed using grid search, followed by model evaluation through 5-fold cross-validation to ensure consistency. Key metrics for assessing classification performance included accuracy for stimulus intensity decoding and F1-macro score for brain area classification, along with precision, recall, and ROC AUC for comprehensive analysis. For binary classification tasks (e.g., low vs. high stimulus intensity), precision and recall were computed using binary averaging, while for multiclass classification tasks (e.g., brain regions), these metrics were calculated with macro averaging. Multiclass ROC AUC was evaluated using a One vs. One approach.

The results provided insights into each model’s effectiveness in distinguishing patterns based on Θ.

## 3 Results

This section presents the main findings from our analysis, focusing on the efficacy of statistical models in approximating neural responses and their application in decoding tasks. Our first objective was to evaluate various combinations of Gaussian and inverse Gaussian distributions (see Table 1) to approximate the Peri-Stimulus Time Histogram (PSTH). Model fitting was validated using the Kolmogorov-Smirnov (KS) test, with the results demonstrating that a three-Gaussian model achieved the best fit based, among all the mixture models considered, on KS test statistics and p-values.

Subsequently, we applied these models to two key decoding tasks: stimulus intensity decoding and brain area classification. For both tasks, performance metrics and feature importance were analyzed. The results show that the model parameters alone, the distribution weights and distribution parameters (means and standard deviations of the Gaussian components), provided sufficient information for accurate decoding, independently of the specific machine learning model applied. Additionally, feature importance analysis revealed which specific Gaussian and which specific components held the greatest relevance for decoding. In the brain area classification task, the parameters of the first Gaussian component were identified as the most significant, indicating that these parameters are particularly informative in distinguishing brain regions. Conversely, in the stimulus intensity decoding task, the parameters of the second Gaussian component showed higher importance, albeit to a lesser extent. Overall, these findings demonstrate that our model parameters are sufficient for high-accuracy decoding of both stimulus intensity and brain area, regardless of the specific machine learning model applied.

### 3.1 Neural Response Data Visualization and Selection

In this section we report the number of neurons considered after applying the selection criteria and we start to compare the response patterns to different stimulation intensities of the two cortex areas.

The database provided by Claar et al. [21] was analyzed to examine the population of neurons for each subject. For the purpose of this study, regular spiking (RS) neurons located within the MO cortex, SS cortex, and sensorimotor-related thalamic nuclei (SM-TH) were considered. To accurately represent the response dynamics following electrical stimulation, an optimal time window was determined through investigation. This analysis identified a significant time window of 300 ms post-stimulus, effectively capturing the relevant neural activity and response patterns. This choice aligns with the findings of Claar et al.’s study, which categorizes the response into three temporal windows following the stimulus: 2-25 ms (early response), 25-150 ms (off period), and 150-300 ms (late response). Figures 2 and 3 show an example of the data considered. Figure 2 shows raster plots of three different neurons belonging to the three brain areas considered, while Figure 3 shows a raster plot of the same neuron stimulated at two different stimulation intensities.

**Figure 2:**
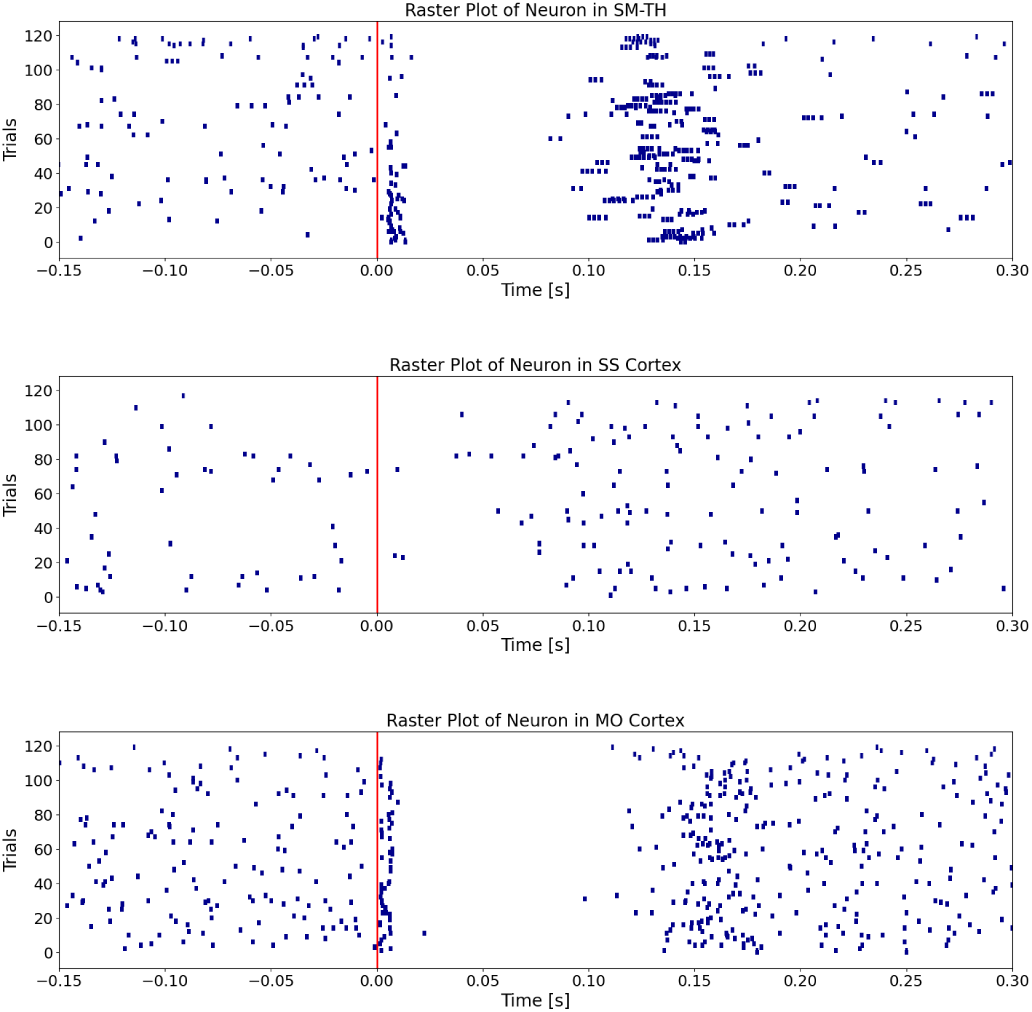
Raster plots of neuronal responses. From top to bottom: a neuron in the SMTH, SS cortex, and MO cortex. The red vertical line indicates stimulus onset. The subject received stimulation in the MO cortex at the highest intensity.

**Figure 3:**
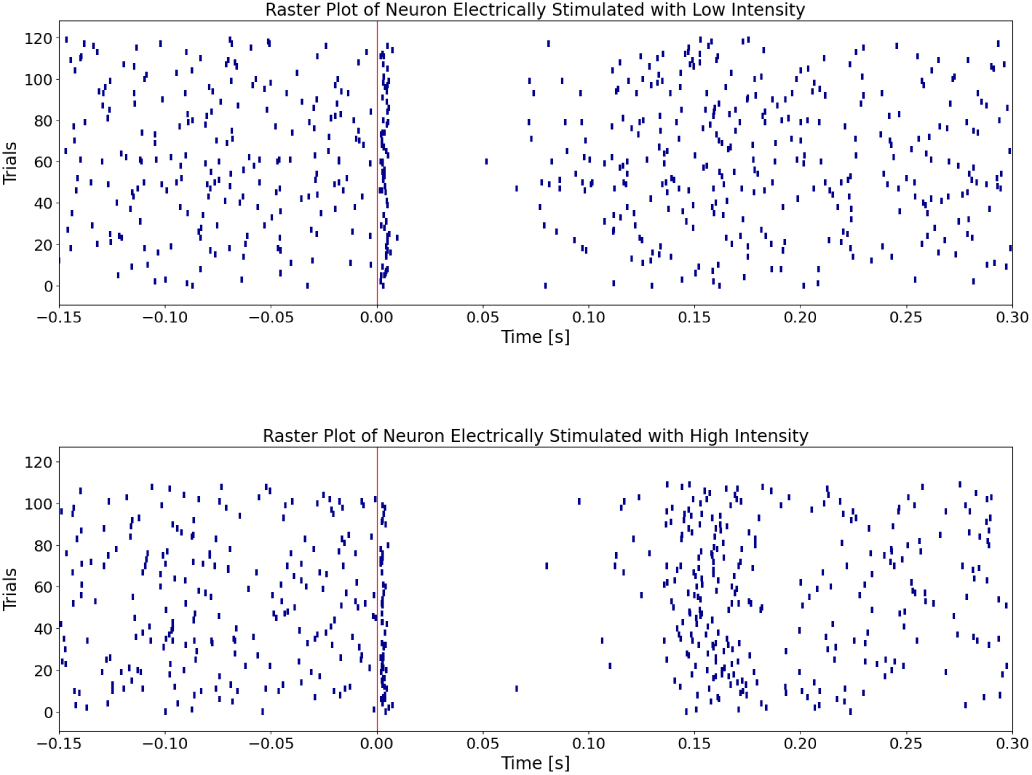
Raster plots of neuronal responses of a MO cortex neuron to low intensity current stimulation (top) and to high intensity current stimulation (bottom). The red vertical line indicates stimulus onset.

After applying the selection criteria described in Section 2.2, we obtained the number of neurons shown in Table 3. For all analyses comparing responses to low and high stimulus intensities, only neurons that responded to both stimulus intensities were considered.

**Table 3:**
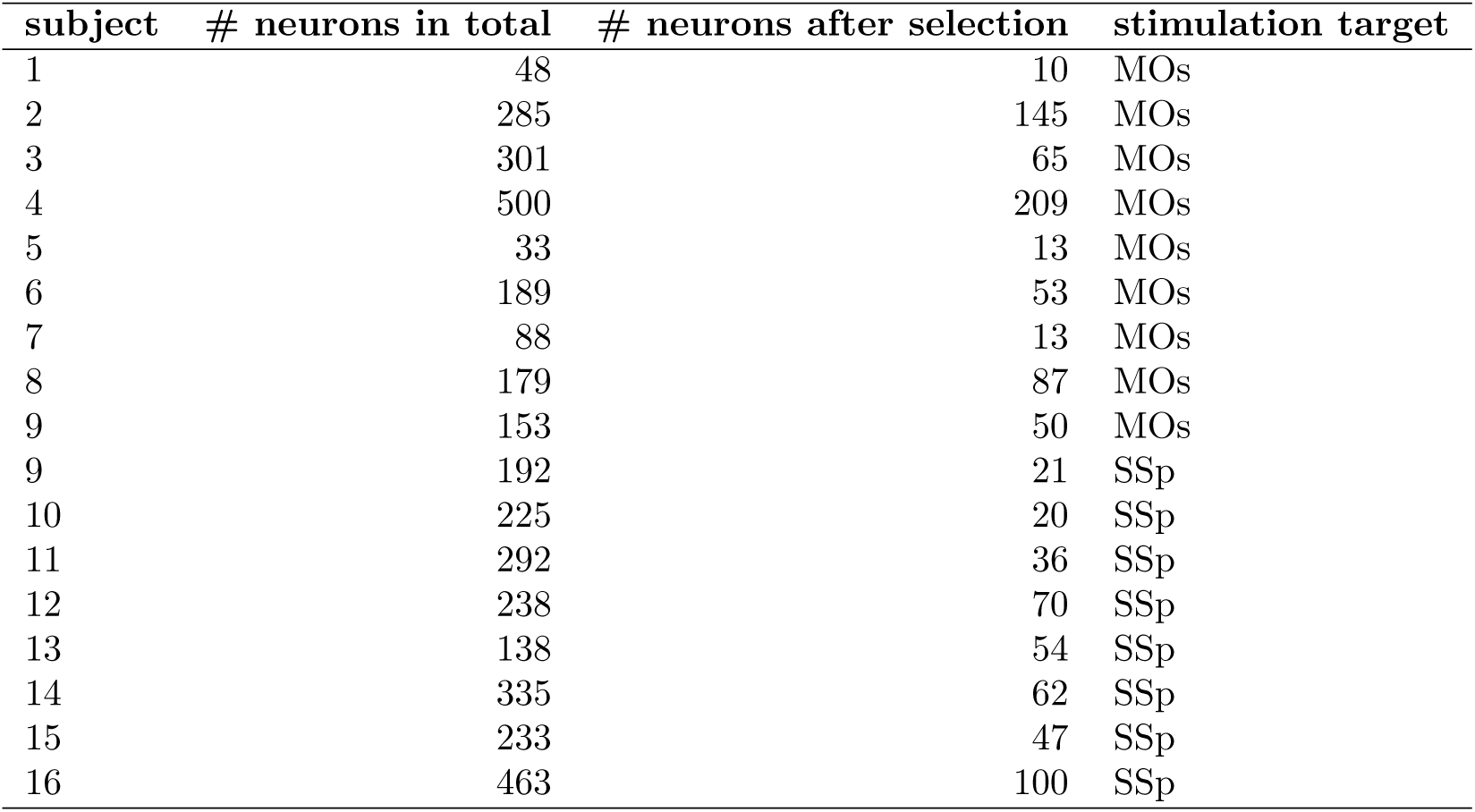
The table below presents the sorted neurons from mouse subjects. It displays the number of neurons identified by Claar et al. [22] during the spike sorting procedure, the number of neurons considered in this study, and the brain region targeted for electrical stimulation.

| subject | # neurons in total | # neurons after selection | stimulation target |
| --- | --- | --- | --- |
| 1 | 48 | 10 | MOs |
| 2 | 285 | 145 | MOs |
| 3 | 301 | 65 | MOs |
| 4 | 500 | 209 | MOs |
| 5 | 33 | 13 | MOs |
| 6 | 189 | 53 | MOs |
| 7 | 88 | 13 | MOs |
| 8 | 179 | 87 | MOs |
| 9 | 153 | 50 | MOs |
| 9 | 192 | 21 | SSp |
| 10 | 225 | 20 | SSp |
| 11 | 292 | 36 | SSp |
| 12 | 238 | 70 | SSp |
| 13 | 138 | 54 | SSp |
| 14 | 335 | 62 | SSp |
| 15 | 233 | 47 | SSp |
| 16 | 463 | 100 | SSp |

Furthermore, by applying the threshold method, we can count the number of bins of PSTHs that exhibit marked response in early and late phases. If there is at least one bin that exceeds the threshold, there is a response (’response’), conversely there is no response (’no response’). When the MO cortex is stimulated, a smaller percentage of the considered neurons exhibit a pronounced early response to low stimulation compared to high stimulation (66.7% and 92.4% respectively) (Table 4). In contrast, the SS cortex does not show a significant difference in early response between low and high stimulation intensities, with a similar percentage (86.4% and 81% respectively) of neurons responding prominently in both cases (Table 5).

**Table 4:** Counts of early and late responses in MO Cortex neurons by stimulation intensity. 66.7% of the population presents a pronounced early response for low stimulation, while 92.4% exhibit this response to high stimulation. 39.4% of the population presents a pronounced late response for low stimulation, while 24.2% exhibit this response to high stimulation.

| Response type | Early phase |  | Late Phase |  |
| --- | --- | --- | --- | --- |
|  | Low stim | High stim | Low stim | High stim |
| No response | 22 | 5 | 40 | 50 |
| Response | 44 | 61 | 26 | 16 |

**Table 5:**
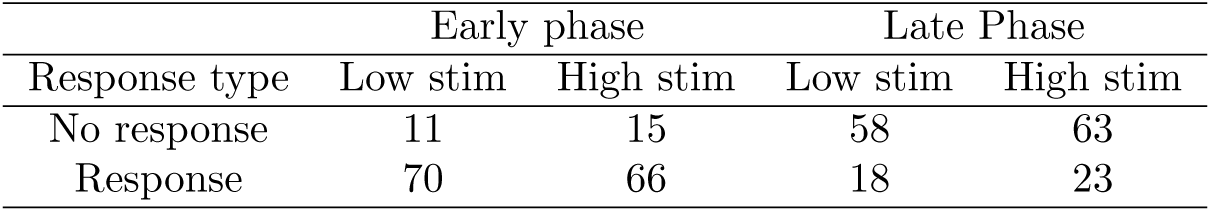
Counts of early and late responses in SS cortex neurons by stimulation intensity. 86.4% of the population presents a pronounced early response for low stimulation, while 81% exhibit this response to high stimulation. 22.2% of the population presents a pronounced late response for low stimulation, while 28.4% exhibit this response to high stimulation.

| Response type | Early phase |  | Late Phase |  |
| --- | --- | --- | --- | --- |
|  | Low stim | High stim | Low stim | High stim |
| No response | 11 | 15 | 58 | 63 |
| Response | 70 | 66 | 18 | 23 |

### 3.2 Model Fitting Analysis and Selection

In this section, we analyze the effectiveness of various statistical models in approximating the PSTH response, focusing on the performance differences between models with different combinations of Gaussian and inverse Gaussian components. We validate the model fits using the KS test.

A model was calculated for each selected neuron (see subSection 2.2), considering factors such as stimulation type (low or high intensity) and stimulation location (i.e. motor (MO) cortex or somatosensory (SO) cortex). For each condition, we tested all relevant combinations of Gaussian and inverse Gaussian distributions (see Table 1), then each model was statistically evaluated as described in subsection 2.5.

Table 6 provides an overview of the tested models, showing their performance metrics. Models were evaluated based on KS distance and p-value, with lower KS distances and higher p-values indicating a better fit to the observed data. To compare the models, we analyzed box plots of the KS statistic and p-value for each model (see Figure 4). As shown, the three-Gaussian model achieves lower KS statistic values and higher p-values, indicating a superior fit to the observed response patterns. This model captures the bi-phasic nature of the neural response more effectively than the other configurations.

**Figure 4:**
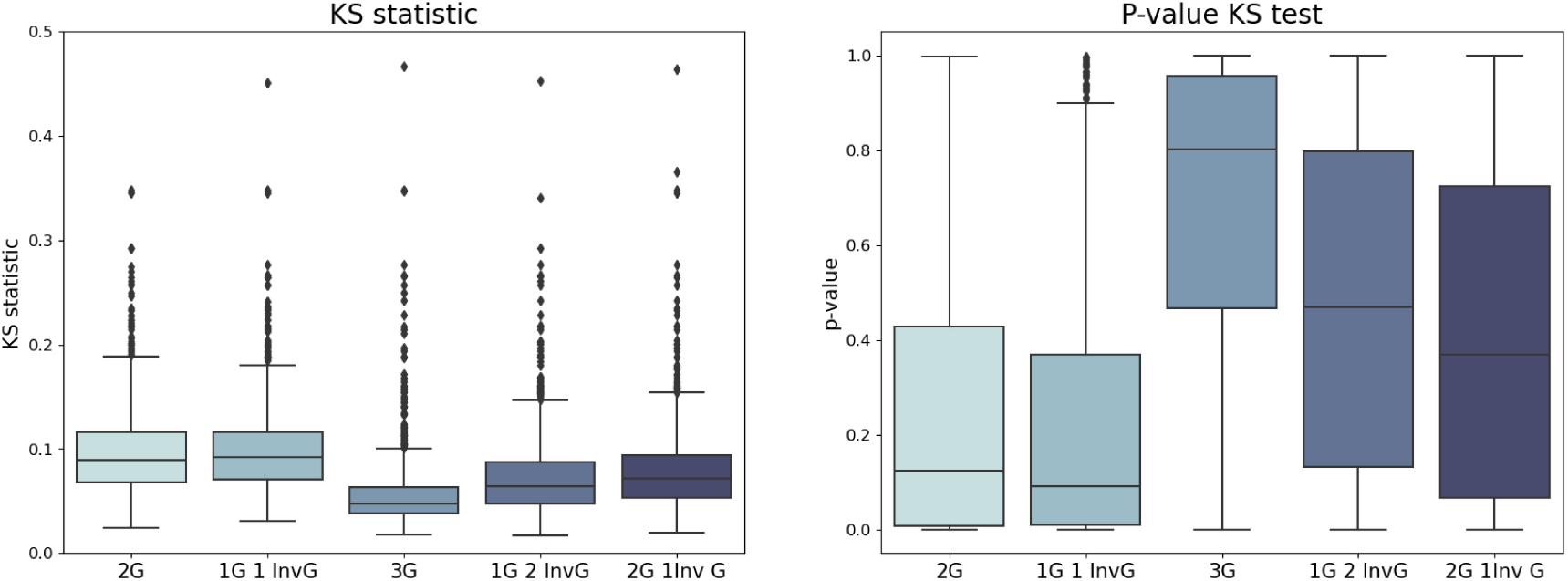
KS Distance and p-value for goodness of fit of different models applied to the mice dataset, modeling the response to the highest stimulation intensity. See Table 6 for model descriptions.

**Table 6:** Summary of the tested models for mice neurons. G stands for Gaussian and InvG stands for Inverse Gaussian. The table displays the configurations and performance metrics (Q1, Q2 and Q3 of KS distance and p-value) of each model.

| Model | $Q1_{dist}$ | $Q2_{dist}$ | $Q3_{dist}$ | $Q1_{pvalue}$ | $Q2_{pvalue}$ | $Q3_{pvalue}$ |
| --- | --- | --- | --- | --- | --- | --- |
| 2G | 0.068200 | 0.089139 | 0.116484 | 0.007642 | 0.123986 | 0.427470 |
| 1G 1 InvG | 0.071173 | 0.091914 | 0.116498 | 0.009290 | 0.091725 | 0.368534 |
| 3G | 0.038127 | 0.047541 | 0.063224 | 0.466125 | 0.800929 | 0.957497 |
| 1G 2InvG | 0.047635 | 0.064307 | 0.087269 | 0.132520 | 0.468810 | 0.798371 |
| 2G 1InvG | 0.053322 | 0.071377 | 0.093844 | 0.067539 | 0.368191 | 0.724447 |

The results above suggest that the three-Gaussian model is the most effective approximation of *f* (*t*) based on the KS test, balancing both KS distance and p-value. Specifically, this model includes one Gaussian component for the early response and two for the late response, capturing both phases of activity. The resulting model is expressed as follows:

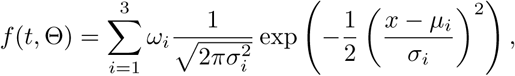

where *ω_i_*represent the weights of *i*-th component, while *µ_i_*and *σ_i_*correspond to the mean and standard deviation of the *i*-th Gaussian component. These parameters were determined by solving the optimization problem in equation (2) and their distribution is discussed in the next subsection. For clarity, the prior distributions of each model parameter are repeated in Table 7 along with their role in describing the response pattern. The response exhibits a triphasic pattern, consisting of an early response (0-50 ms) and a late response (50-300 ms), separated by an off period. The first Gaussian captures the early response, while the second and third Gaussians represent the late response. As a result, the late response can be divided into two distinct components, with the first of these—within the scope of this study—being the one that contains more relevant information about the response dynamics.

**Table 7:** Model parameters, their prior distributions, roles in the model, and physiological meanings. *U* refers to the Uniform distribution.

| Model Parameter | Prior Distribution | Role in the model | Physiological interpretation |
| --- | --- | --- | --- |
| $\mu_1$ | $\sim U(0, 0.05)$ | mean of the first Gaussian | mean latency of the early response |
| $\sigma_1$ | $\sim U(10^{-6}, 1)$ | standard deviation of the first Gaussian | variability of the early response |
| $\mu_2$ | $\sim U(0.05, 0.3)$ | mean of the second Gaussian | mean latency of the first part of late response (i.e. rebound excitation) |
| $\sigma_2$ | $\sim U(10^{-6}, 1)$ | standard deviation of the second Gaussian | variability of the first part of late response (i.e. rebound excitation) |
| $\mu_3$ | $\sim U(\mu_2, 0.3)$ | mean of the third Gaussian | mean latency of the second part of late response |
| $\sigma_3$ | $\sim U(10^{-6}, 1)$ | standard deviation of the third Gaussian | variability of the second part of late response |
| $[\omega_1, \omega_2, \omega_3]$ | $\sim Dir(1, 1, 1)$ | weights associated to the Dirichlet mixture model | influence of each component in explaining the observed data distribution. |

For further illustration, Figures 5, 6 and 7 compare observed and model-predicted CDFs across different brain regions neurons. These visualizations highlight the model’s fit for individual neurons, with the three-Gaussian model showing the closest approximation.

**Figure 5:**
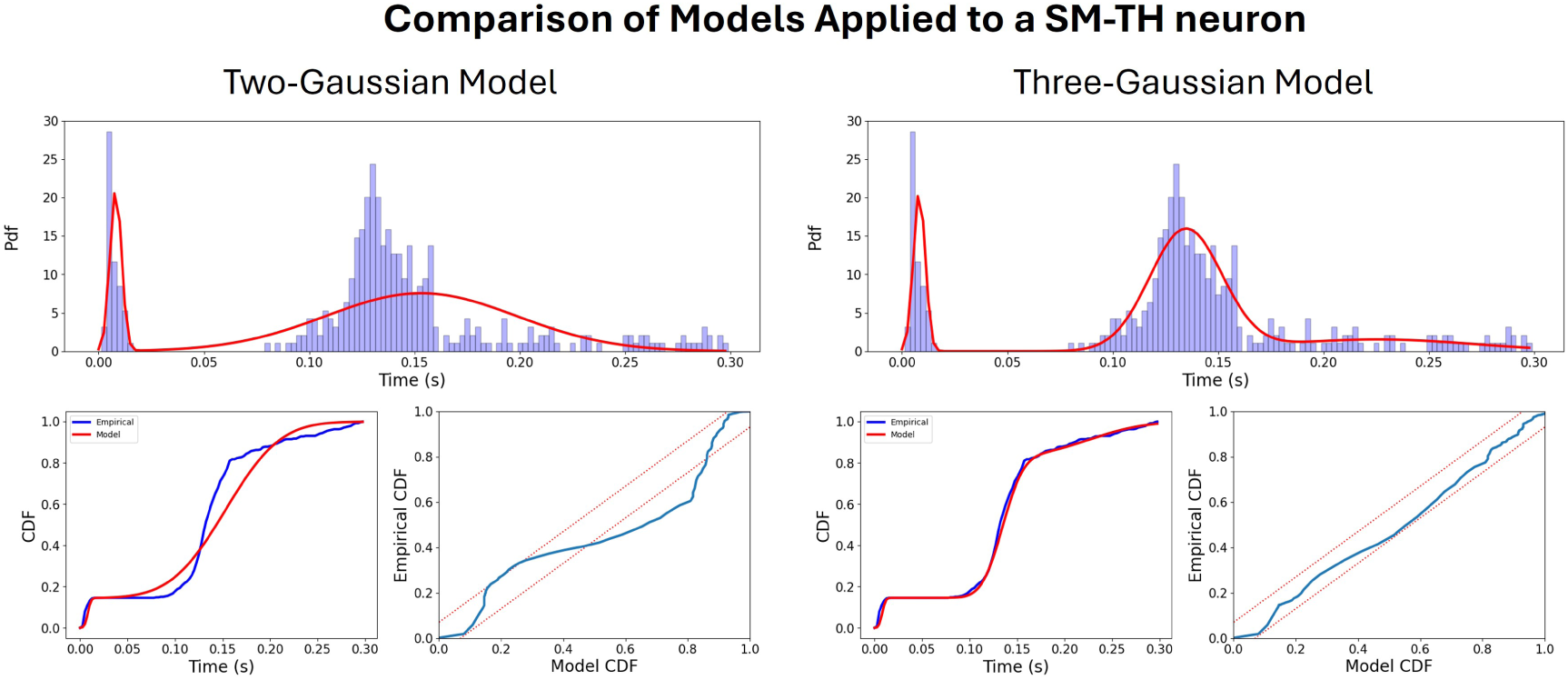
Comparison between the Two-Gaussian Model (left) and the Three-Gaussian Model (right) applied to a Neuron from the sensorimotor-related thalamic nuclei (SM-TH). The KS test results for the Two-Gaussian model indicate a KS distance of 0.19 with a p-value less than 0.001. In comparison, the Three-Gaussian model shows a KS distance of 0.04 with a p-value of 0.66.

**Figure 6:**
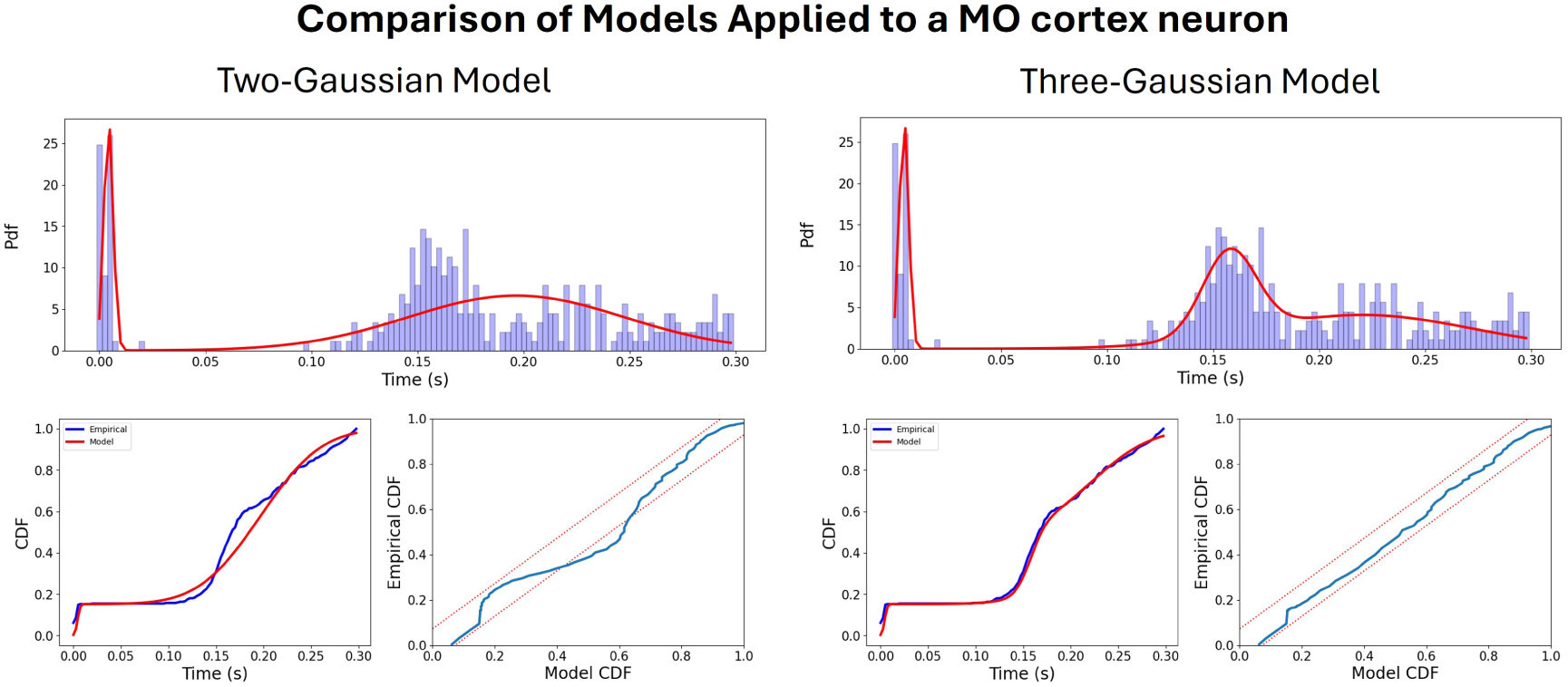
Comparison between the Two-Gaussian Model (left) and the Three-Gaussian Model (right) applied to a Neuron from the MO cortex. The KS test results for the Two-Gaussian model indicate a KS distance of 0.12 with a p-value less than 0.001. In comparison, the Three-Gaussian model shows a KS distance of 0.03 with a p-value of 0.82.

**Figure 7:**
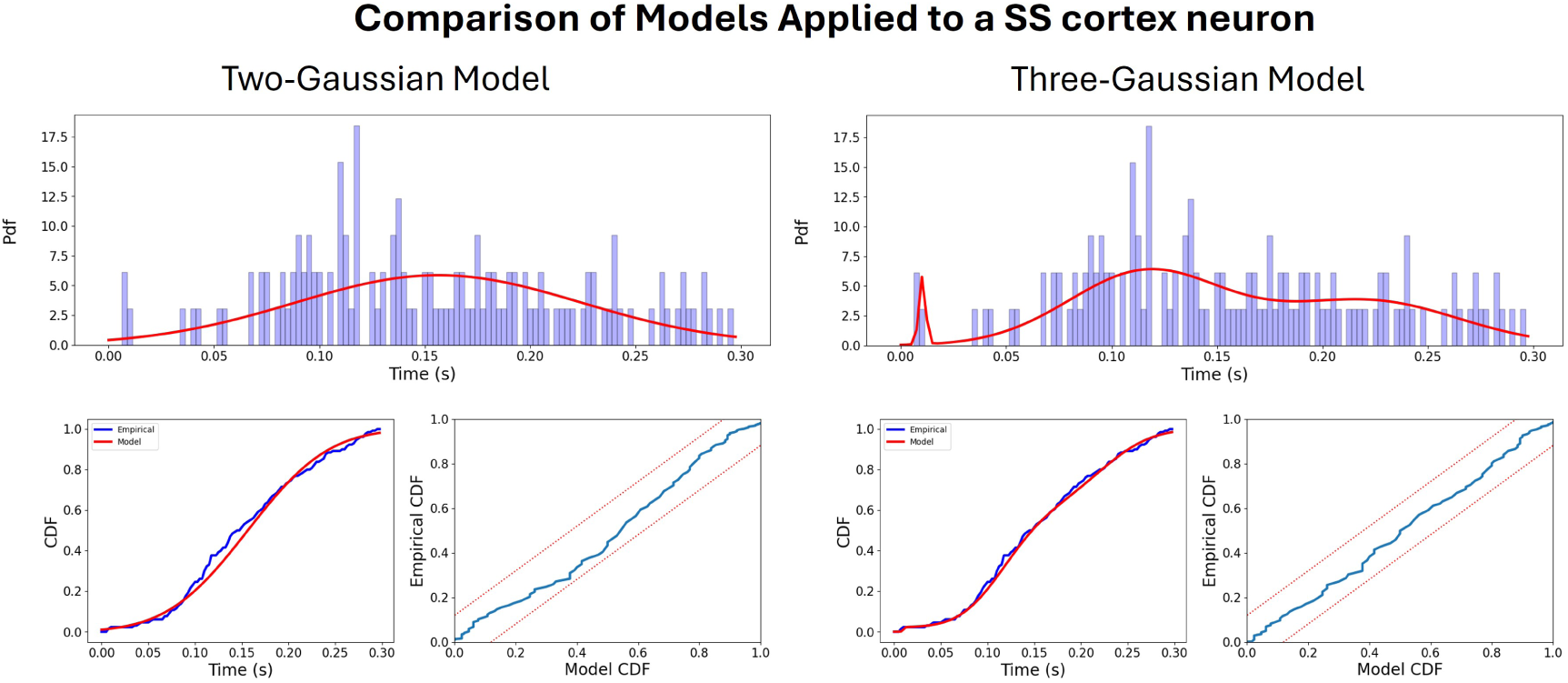
Comparison between the Two-Gaussian Model (left) and the Three-Gaussian Model (right) applied to a Neuron from the SS cortex. The KS test results for the Two-Gaussian model indicate a KS distance of 0.08 with a p-value of 0.36. In comparison, the Three-Gaussian model shows a KS distance of 0.04 with a p-value of 0.96.

### 3.3 Model Parameters Distribution Visualization

In this section, various plots related to the parameters (*µ_i_, σ_i_*) representing the mean, and standard deviation of the *i*-th Gaussian used in the model are presented and their characteristics discussed in order to provide a comprehensive understanding of the neural response dynamics under different stimulation conditions. These parameters are used to elucidate how different brain zones (MO cortex, SS cortex, and SM-TH) respond to stimuli in terms of the timing and variability of their neural activity. For more information about the model parameters see Tables 12, 13, 14, 15 in Appendix.

Figures 8, 10, 12, and 13 illustrate the differences between the distribution of the model parameters (*µ_i_, σ_i_*), based on stimulation intensity (low vs. high) or the specific brain regions to which the neurons belong (MO cortex, SS cortex, and SM-TH). The statistical analysis results highlight significant variations among the different brain regions and stimulation intensities. The values in the text are presented as the median and IQR (inter-quartile range: 25th and 75th percentiles). To ensure comparable stimulation intensities applied to the MO cortex and the SS cortex, we conducted Mann-Whitney tests on the intensities applied to each subject in the study. The results revealed no significant statistical difference between the stimulation intensities across cortical areas (see Figure 21).

**Figure 8:**
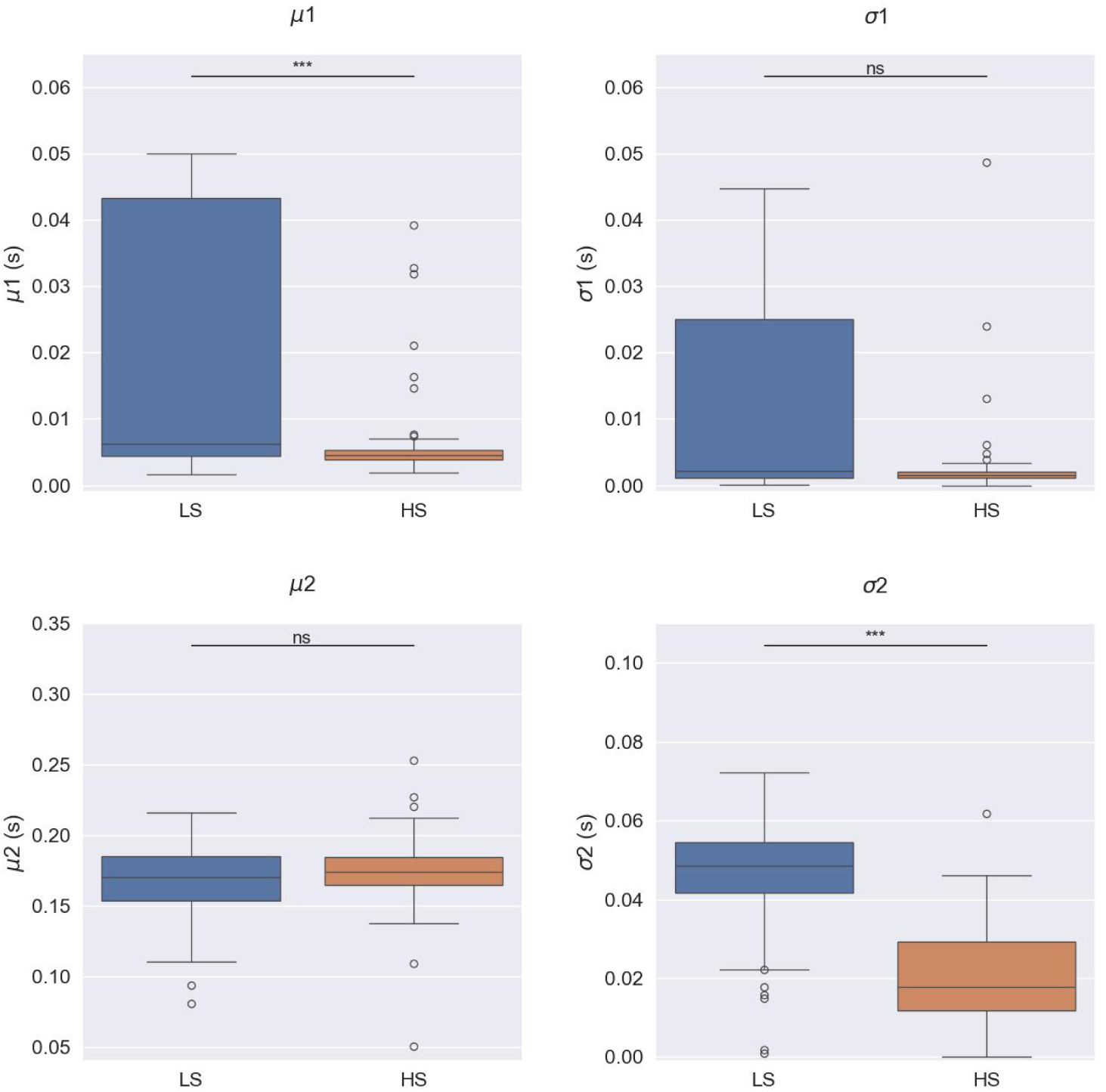
Boxplot of model parameters for MO cortex neurons when electrically stimulated with *low* and *high* current intensities. Boxplots show median, 25th, and 75th percentiles; whiskers extend from the box by 1.5× the IQR. Wilcoxon signed-rank tests: n.s. no significant evidence to reject null hypothesis (*p >* 0.05), * weak evidence to reject null hypothesis (0.01 *< p <* 0.05), ** strong evidence to reject null hypothesis (0.001 *< p <* 0.01), and *** very strong evidence to reject null hypothesis (*p <* 0.001). In the early response, the parameter *µ*_1_ (*p <* 0.001) is higher and more variable at low intensities compared to high intensities. In the late response, *σ*_2_ (*p <* 0.001) is higher for low intensities compared to high intensities. For rebound excitation, comparable mean latencies are observed across intensities, but higher intensities result in narrower response curves.

Figures 8 and 10 display the parameter distributions when varying stimulus intensities are used to directly trigger the MO and SS cortex. By looking at Figure 8, we can observe that for the MO cortex stimulation dataset, the early response shows that *µ*_1_ and *σ*_1_ present higher and more variable values for low intensities (0.006 [0.004 - 0.043] s and 0.002 [0.001 - 0.025] s) compared to higher intensities (0.005 [0.004 - 0.005] s and 0.001 [0.001 - 0.002] s). This suggests that higher intensities elicit a more defined early response, whereas lower intensities lead to greater variability. In the late response, there is no significant visible variation in *µ*_2_ across the two intensities (0.170 [0.154 - 0.185] s and 0.174 [0.165 - 0.185] s), while *σ*_2_ tends to be higher for low intensity (0.049 [0.042 - 0.055] s) with respect to high intensity (0.018 [0.012 - 0.029] s). The rebound excitation occurs at comparable mean latencies between the two intensities; however, higher intensities may result in narrower curves, which leads to a more precise response. Figure 9 presents an example of electrically evoked responses to low and high intensity stimulation within the same neuron of the MO cortex. A similar pattern is present when the SS cortex is stimulated, but it is less pronounced (see Figure 10). Figure 11 presents an example of electrically evoked responses to low and high intensity stimulation within the same neuron of the SS cortex.

**Figure 9:**
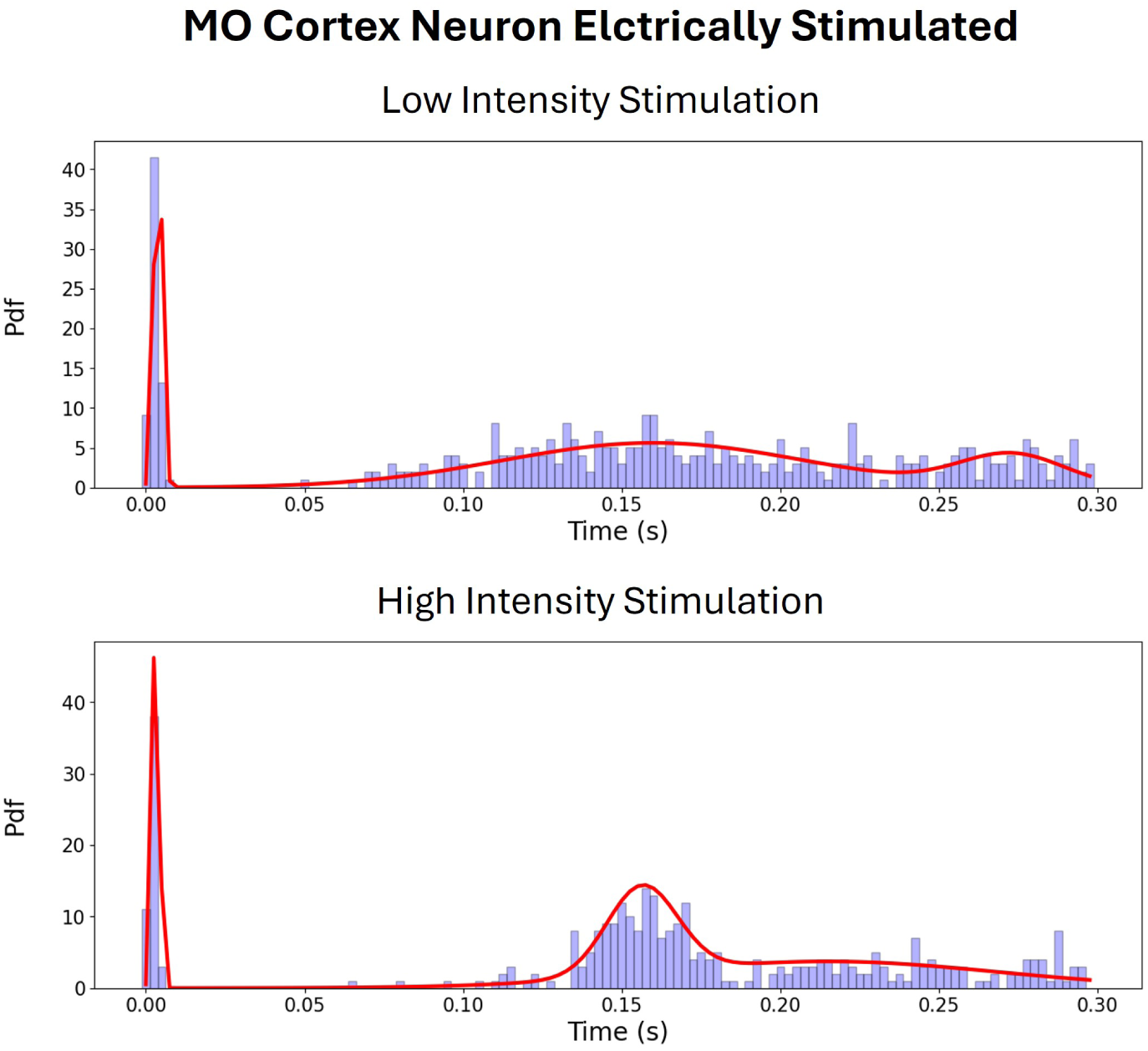
Comparison of electrically evoked responses to low and high intensity stimulation within the same neuron of the MO cortex.

**Figure 10:**
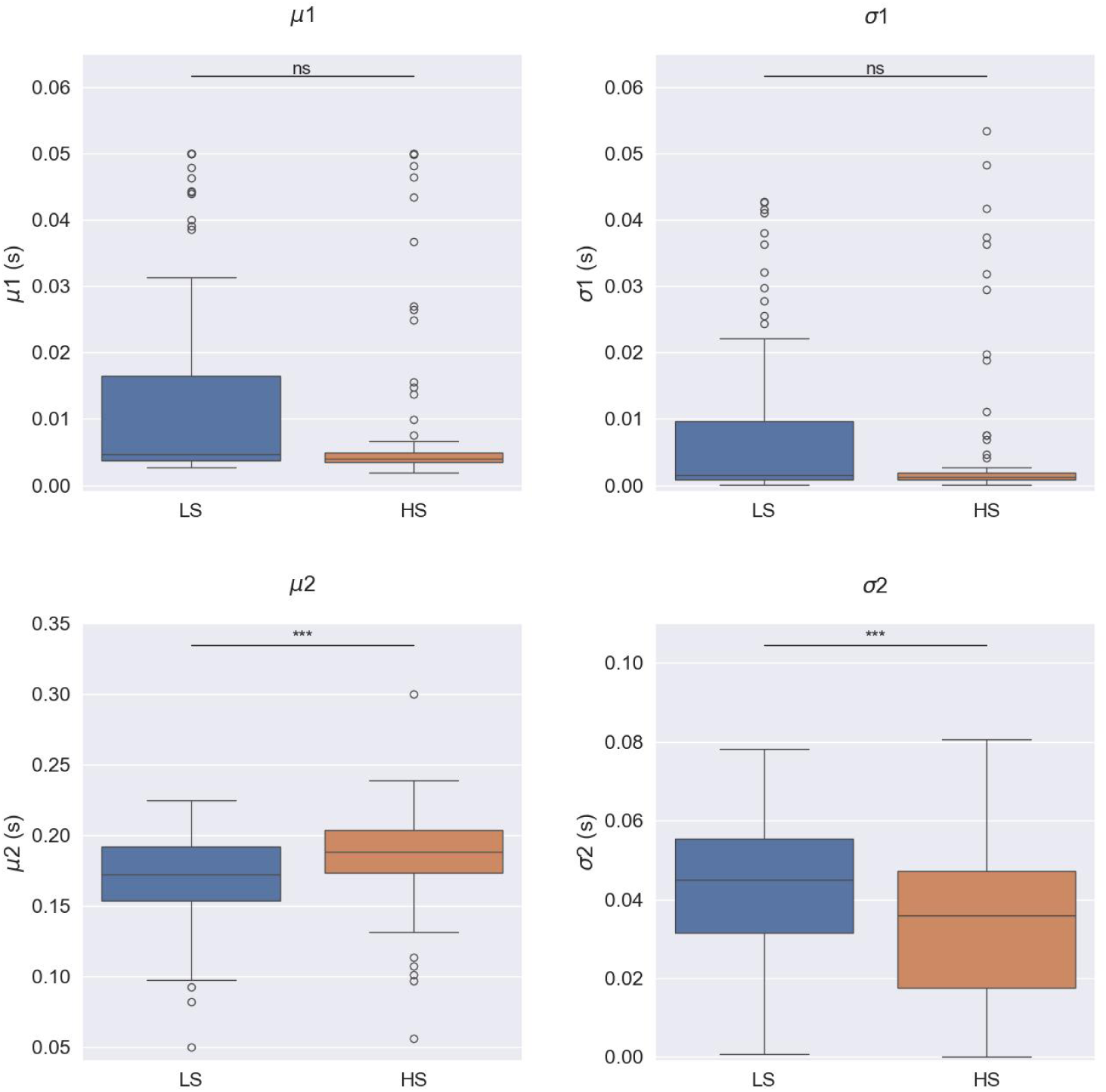
Boxplot of model parameters for SS cortex neurons when electrically stimulated with *low* and *high* current intensities. Boxplots show median, 25th, and 75th percentiles; whiskers extend from the box by 1.5× the IQR. Wilcoxon signed-rank tests: n.s. no significant evidence to reject null hypothesis (*p >* 0.05), * weak evidence to reject null hypothesis (0.01 *< p <* 0.05), ** strong evidence to reject null hypothesis (0.001 *< p <* 0.01), and *** very strong evidence to reject null hypothesis (*p <* 0.001). In the early response, the parameter *µ*_1_ (0.005 *> p*) is higher and more variable at low intensities compared to high intensities. In the late response, *µ*_2_ (*p <* 0.001) presents higher values for high intensities, whereas *σ*_2_ (*p <* 0.001) is higher for low intensities compared to high intensities. Therefore, the rebound excitation occurs at higher latencies at high intensities presenting narrower response curves.

**Figure 11:**
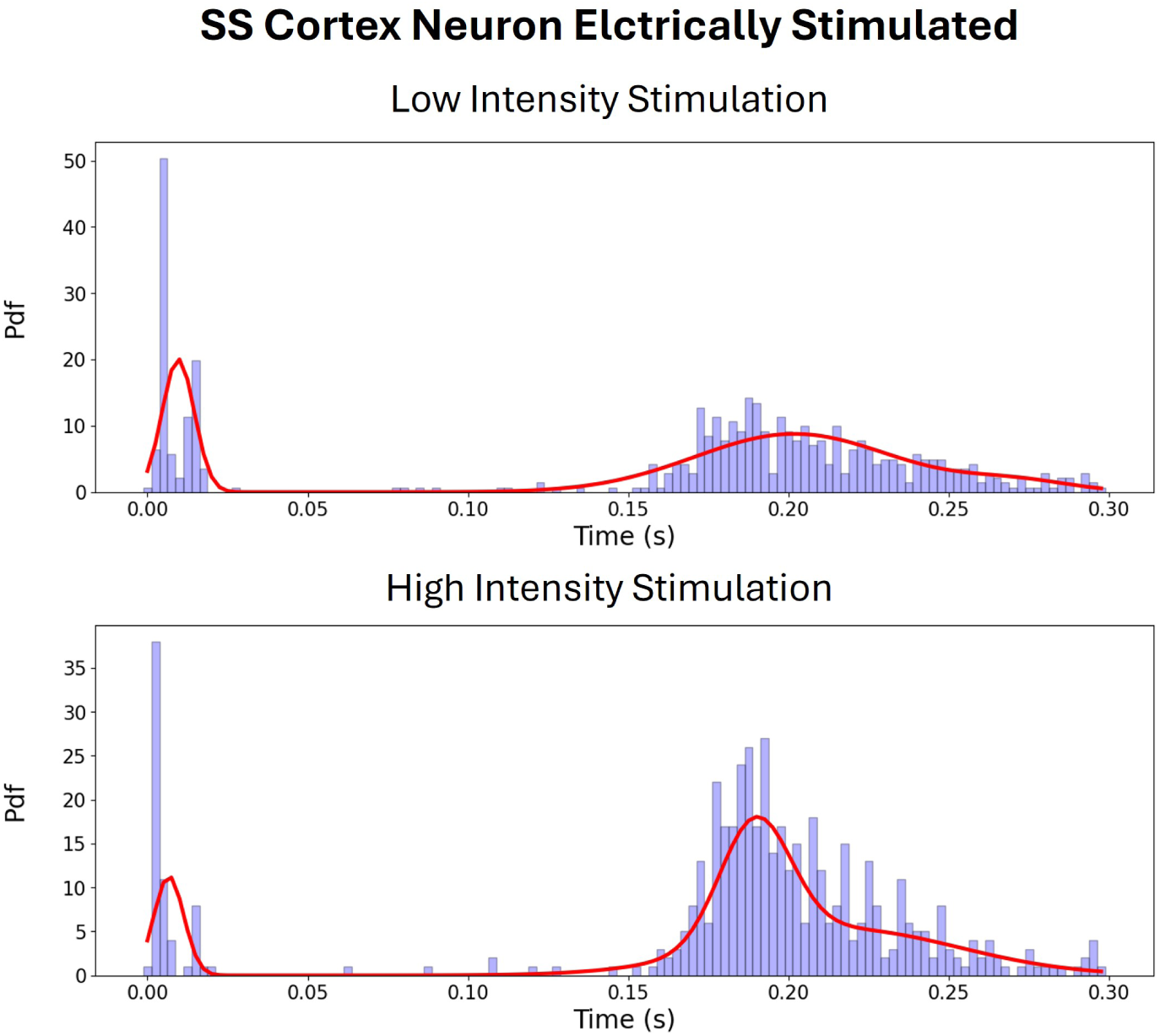
Comparison of electrically evoked responses to low and high intensity stimulation within the same neuron of the SS cortex.

Figure 12 illustrates the differences in model parameters across various brain zones (SS cortex, MO cortex, and SM-TH) when the MO cortex is stimulated with the highest intensity. In contrast, Figure 13 shows these differences when the SS cortex is stimulated with the highest intensity.

**Figure 12:**
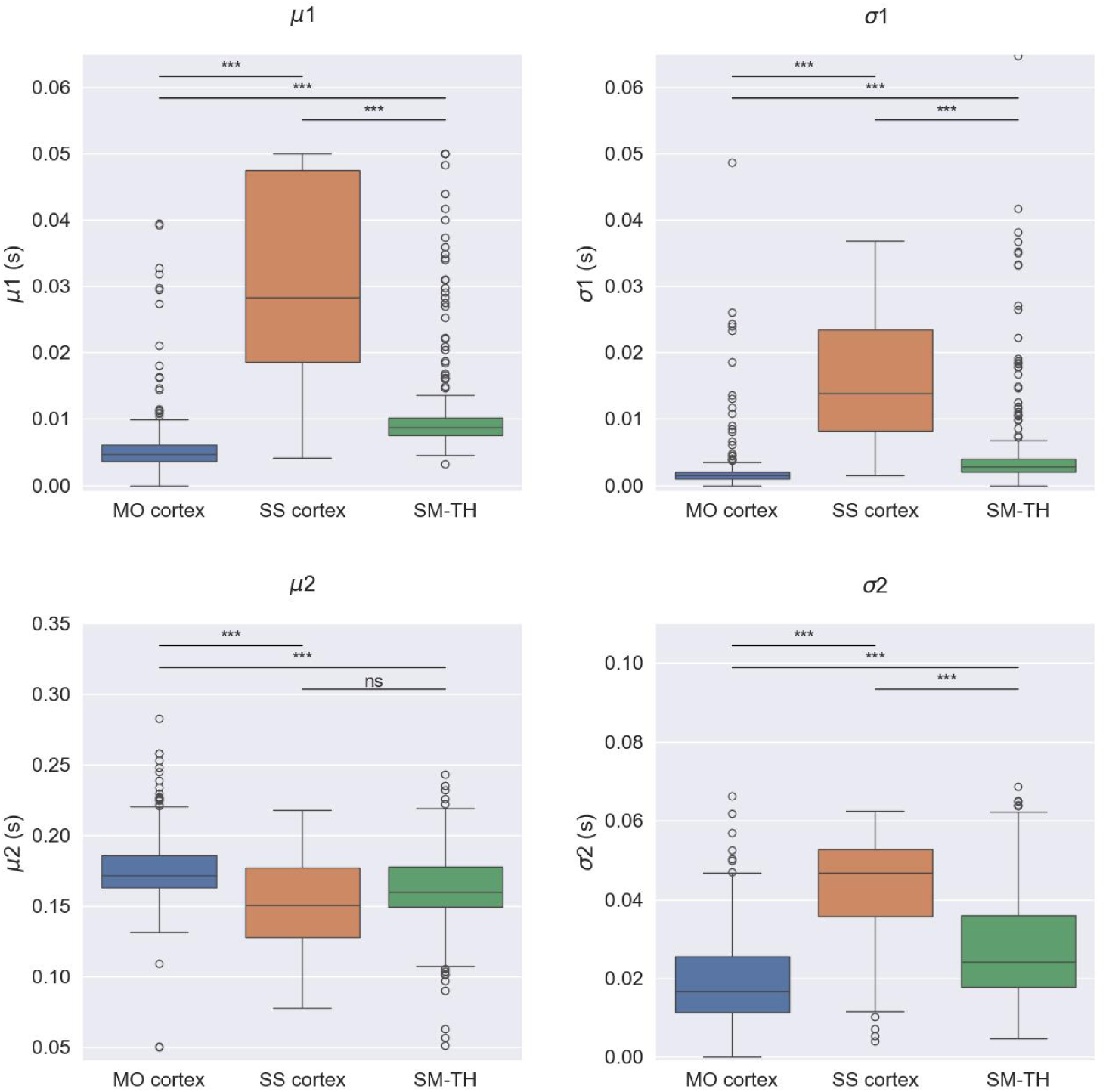
Boxplot of model parameters for SM-TH, MO Cortex, and SS Cortex neurons when MO cortex is electrically stimulated with *high* current intensities. Boxplots show median, 25th, and 75th percentiles; whiskers extend from the box by 1.5 × the IQR. Mann-Whitney tests: n.s. no significant evidence to reject null hypothesis (*p >* 0.05), * weak evidence to reject null hypothesis (0.01 *< p <* 0.05), ** strong evidence to reject null hypothesis (0.001 *< p <* 0.01), and *** very strong evidence to reject null hypothesis (*p <* 0.001). The MO cortex shows the earliest response, followed by the SM-TH and SS cortex, with specific average *µ*_1_ values (*p <* 0.001). Both the MO cortex and SM-TH have sharp early responses, indicated by lower mean *σ*_1_ values (*p <* 0.001), while the SS cortex displays greater variability with higher *σ*_1_ (*p <* 0.001). For the late response, the MO cortex has the longest latency, while the SS cortex and SM-TH initiates earlier, as reflected in their respective *µ*_2_ values (*p <* 0.001). During this phase, the MO cortex and SM-TH demonstrate lower *σ*_2_ values compared to the SS (*p <* 0.001) cortex, indicating less variability.

**Figure 13:**
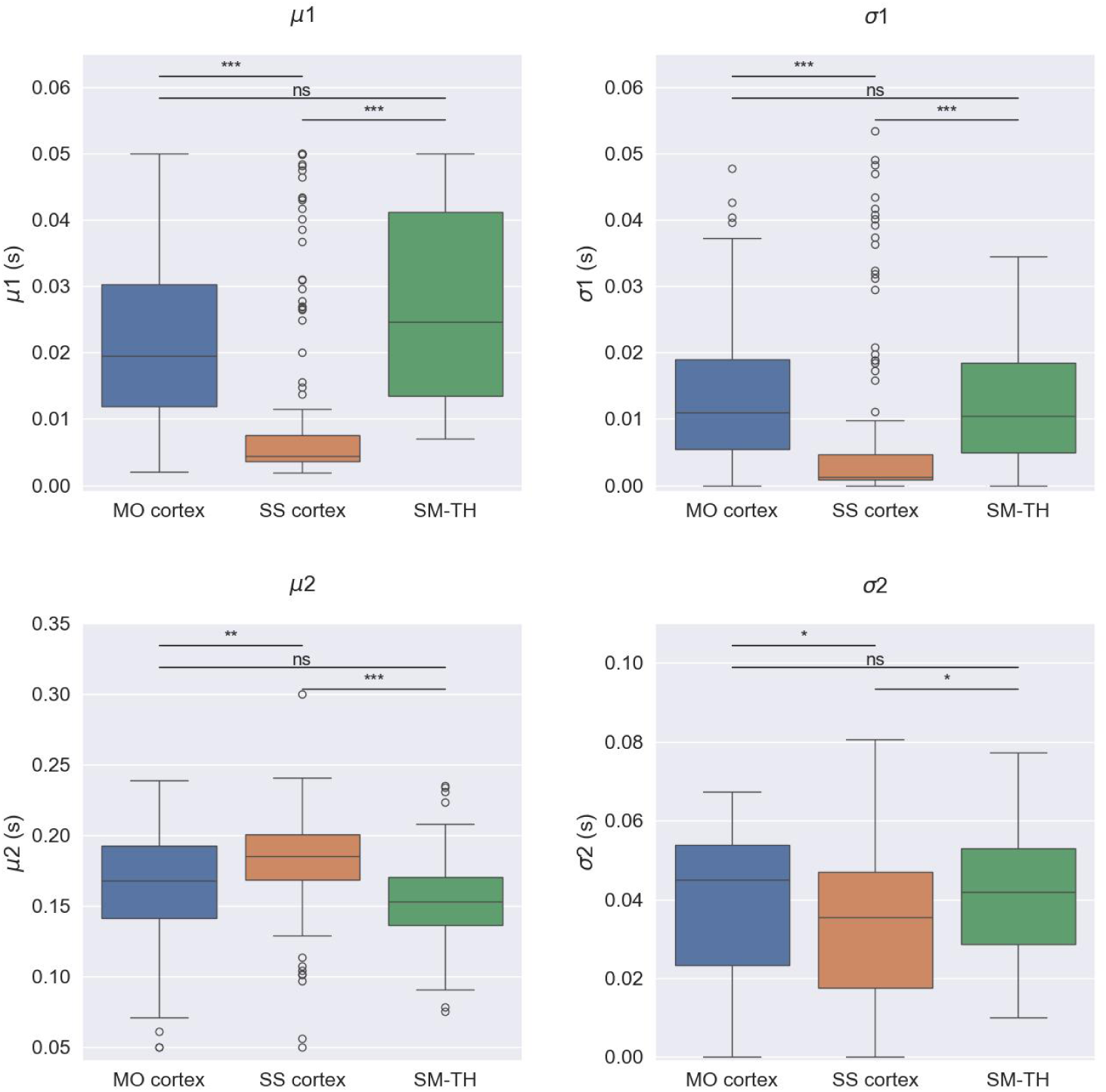
Boxplot of model parameters for SM-TH, MO Cortex, and SS Cortex neurons when SS Cortex is electrically stimulated with *high* current intensities. Boxplots show median, 25th, and 75th percentiles; whiskers extend from the box by 1.5× the IQR. Mann-Whitney tests: n.s. no significant evidence to reject null hypothesis (*p >* 0.05), * weak evidence to reject null hypothesis (0.01 *< p <* 0.05), ** strong evidence to reject null hypothesis (0.001 *< p <* 0.01), and *** very strong evidence to reject null hypothesis (*p <* 0.001). When the SS cortex is stimulated, it exhibits the earliest response, followed by the MO cortex and the SM-TH, as indicated by their median *µ*_1_ values (*p <* 0.05). The SS cortex shows a sharper early response compared to the MO cortex and SM-TH, reflected in its lower median *σ*_1_ values (*p <* 0.001). For the late response, the SS cortex has the longest latency, while the SM-TH and MO cortex initiate the response the earliest, as shown by their respective *µ*_2_ values (0.01 *> p*). During this phase, the not directly stimulated areas show relatively similar *σ*_2_ values, whereas the stimulated cortex shows lower values (*p <* 0.05).

Examining the box plot of *µ*_1_ in Figure 12, we observe that upon MO cortex stimulation, the MO cortex exhibits the earliest response, followed by the SM-TH, and finally the SS cortex (with average *µ*_1_ values of 0.005 [0.004 - 0.006] s, 0.009 [0.008 - 0.010] s, and 0.028 [0.019 - 0.048] s, respectively). Additionally, both the MO cortex and SM-TH display a sharp early response, characterized by a median *σ*_1_ of 0.001 [0.001 - 0.002] s and 0.003 [0.002 - 0.004] s, whereas the SS cortex exhibits a more variable response (median *σ*_1_ of 0.014 [0.008 - 0.024] s). Regarding the late response, the MO cortex demonstrates the greatest latency on average, while the SS cortex initiates the late response the earliest, followed by the SM-TH (median *µ*_2_ values of 0.172 [0.163 - 0.186] s, 0.151 [0.128 - 0.177] s, and 0.160 [0.149 - 0.178] s, respectively). During this phase, the MO cortex and SM-TH exhibit lower values of *σ*_2_ compared to the SS cortex (median *σ*_2_ values of 0.017 [0.011 - 0.026] s, 0.024 [0.018 - 0.036] s, and 0.047 [0.036 - 0.053] s, respectively). In contrast, examining the box plots in Figure 13, when the SS cortex is stimulated, the SS cortex is the first to exhibit an early response, followed by the MO cortex and finally the SM-TH (with median *µ*_1_ values of 0.004 [0.004 - 0.008] s, 0.020 [0.012 - 0.030] s, and 0.025 [0.014 - 0.041] s, respectively). The SS cortex displays a sharper early response compared to the MO cortex and SM-TH (median *σ*_1_ values of 0.001 [0.001 - 0.005] s, 0.011 [0.005 - 0.019] s, and 0.010 [0.005 - 0.019] s, respectively). Regarding the late response, the SS cortex demonstrates the greatest latency on average, while the SM-TH initiates the late response the earliest, followed by the MO cortex (median *µ*_2_ values of 0.185 [0.169 - 0.200] s, 0.153 [0.136 - 0.17] s, and 0.168 [0.142 - 0.192] s, respectively). In this case, the three areas present not so different values of *σ*_2_ (values of 0.035 [0.018 - 0.047] s for SS cortex, 0.042 [0.029 - 0.053] s for SMTH, and 0.045 [0.023 - 0.054] s for MO cortex).

Scatter plots 14 and 15 further detail the distribution of features in the early response of SS and MO cortex neurons, depending on the stimulation site, revealing a symmetrical pattern between the two areas. When the MO cortex is stimulated, the SS cortex presents similar responses to those displayed by the MO cortex when the SS cortex is stimulated. For the stimulated cortex, the early response is sharp (indicating low standard deviations *σ*_1_) and occurs close to the stimulus onset (low values of *µ*_1_). In contrast, the non-stimulated cortex exhibits a bell-shaped response with greater variability (i.e., a wider range of values for both *µ*_1_ and *σ*_1_). The figures illustrate a quasi-linear relationship between the two parameters: as the average latency of the early response (*µ*_1_) increases, so does its variability (*σ*_1_).

An additional analysis was conducted to examine potential parallels between the model’s features and findings from the study by Claar et al. [22]. It was observed that the median first spike latency following stimulus onset was shorter in the stimulated cortex compared to the thalamus during the early response, with the opposite trend noted during the late response. We computed the median values of *µ*_1_ (i.e. mean of early response) in the early response and the median values of *µ*_2_ (i.e. mean of the first part of late response) in the late response for every subject. As shown in Figure 16, our findings mirror those of Claar et al. The temporal order of responses is preserved: in the early response phase, the stimulated cortex exhibits activity before the SM-TH, while in the late response phase, it shows activity after the SM-TH.

**Figure 14:**
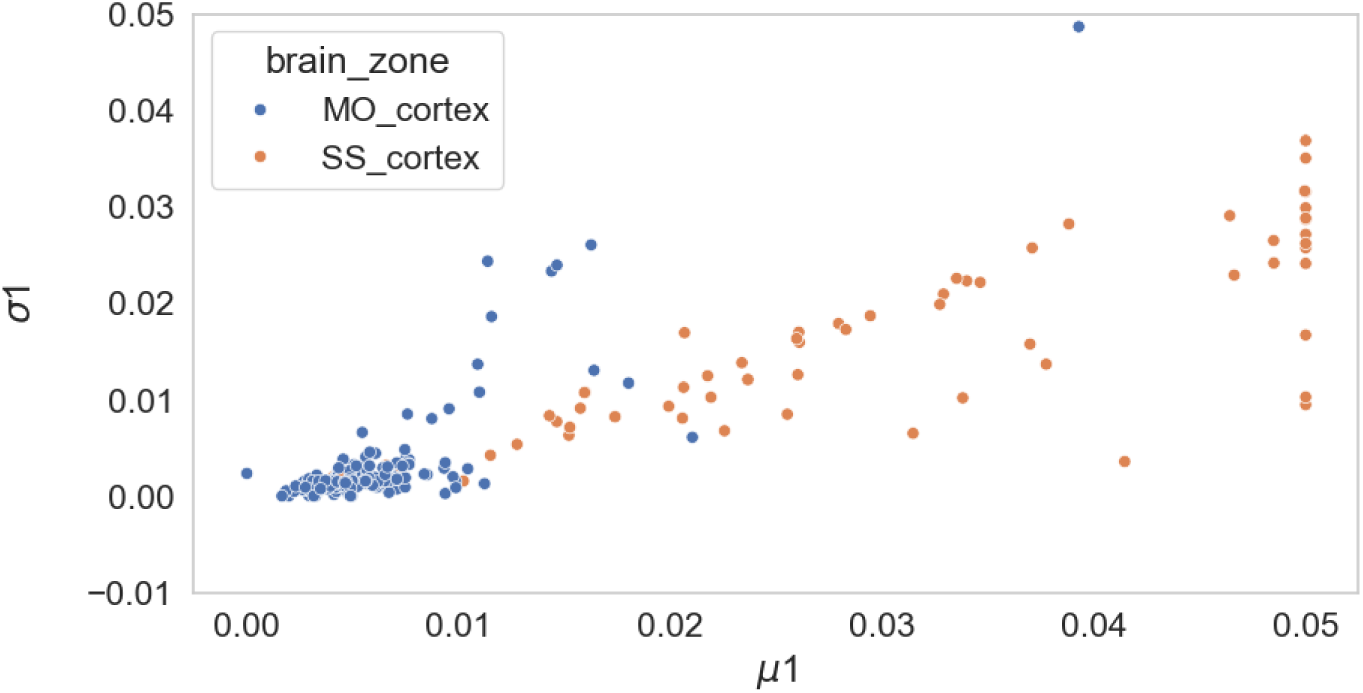
Model parameters (mean *µ*_1_ and standard deviation *σ*_1_ of the first gaussian curve) describing the early response of brain cortex areas when MO cortex is stimulated. Each dot represents a neuron. The image shows a quasi-linear relationship between the two parameters: as the average latency of the early response increases, so does its variability.

**Figure 15:**
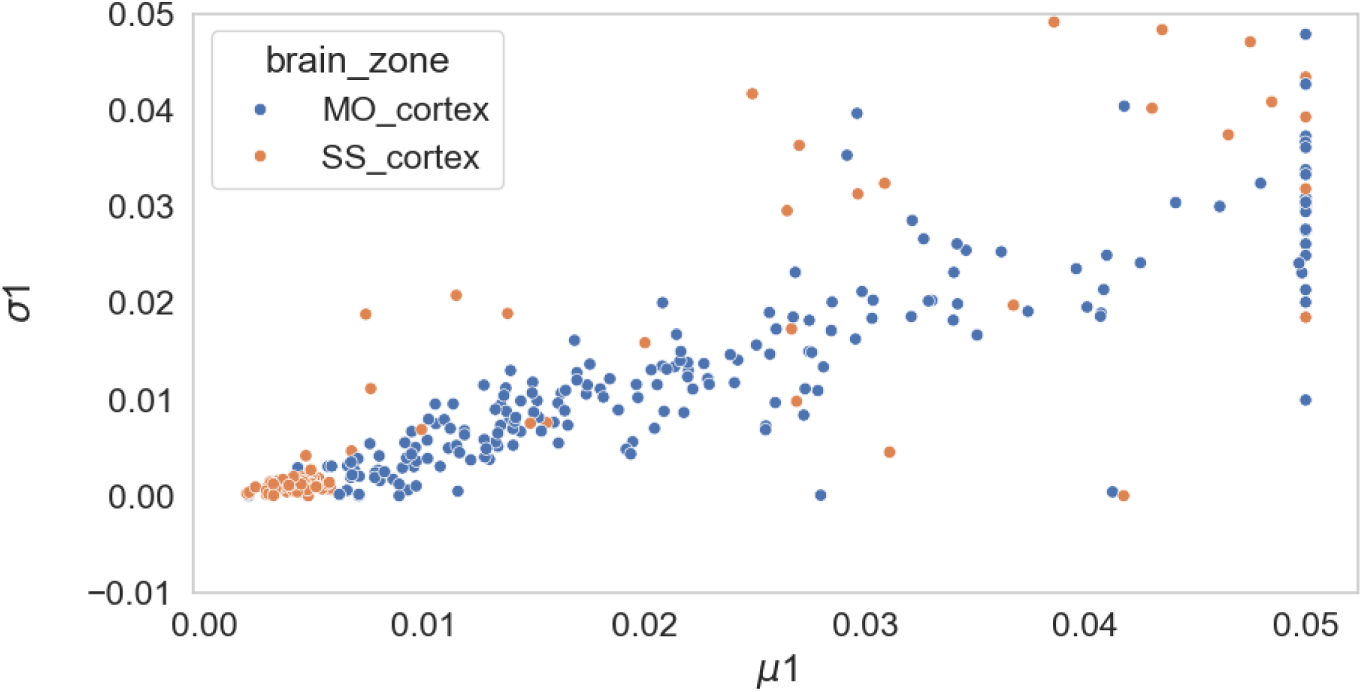
Model parameters (mean *µ*_1_ and standard deviation *σ*_1_ of the first gaussian curve) describing the early response of brain cortex areas when SS cortex is stimulated. Each dot represents a neuron. The image shows a quasi-linear relationship between the two parameters: as the average latency of the early response increases, so does its variability.

**Figure 16:**
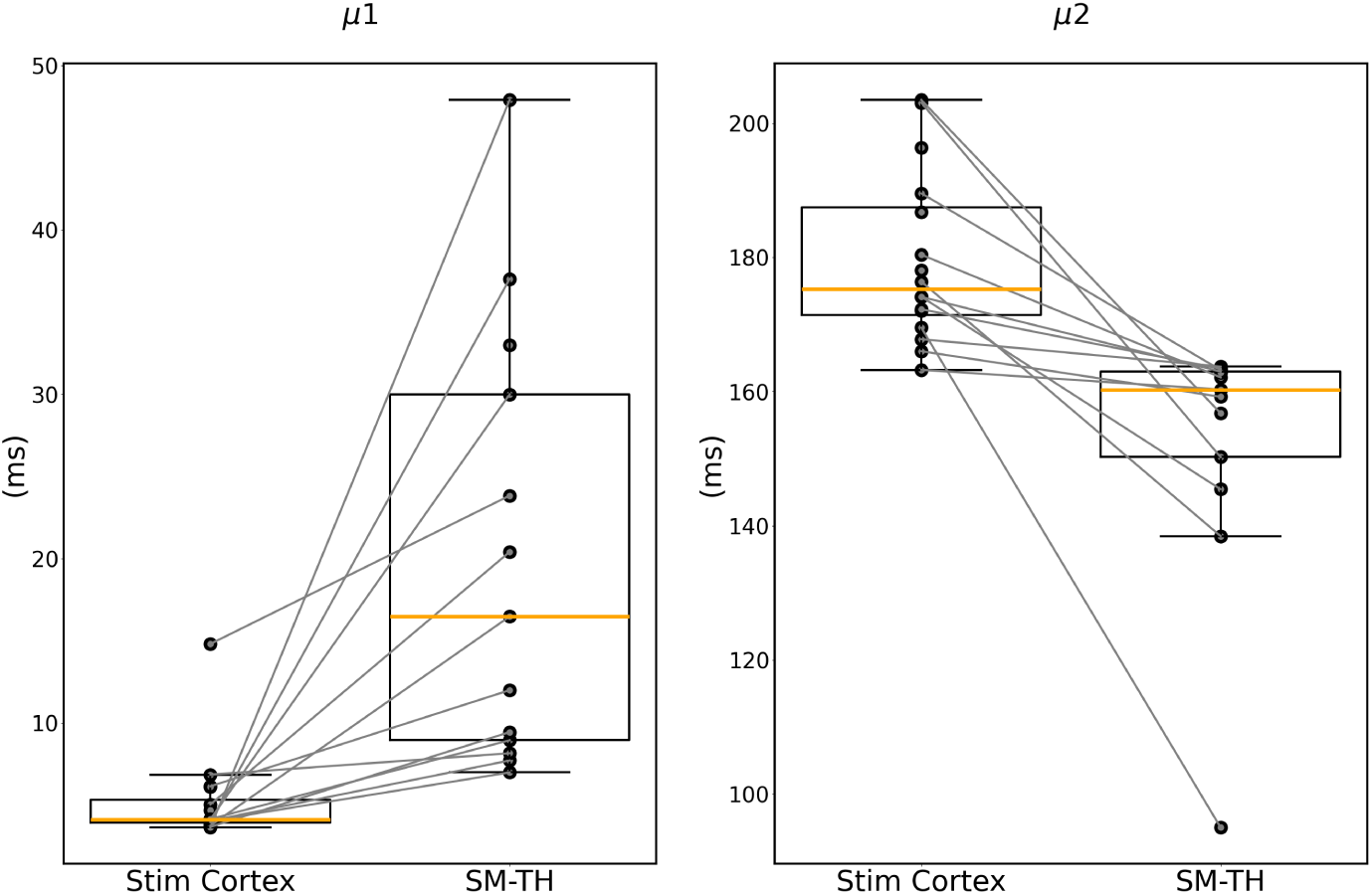
Median values of *µ*_1_ and *µ*_2_ computed for responsive neurons in stimulated cortex (left) and in associated SM-TH (right). Each circle represents the median of *µ*_1_ or *µ*_2_ computed for one subject, represented by the connecting gray lines.

### 3.4 Stimulus Intensity Decoding

We here present the results of decoding stimulation intensities (low vs high) based on neural responses, using binary ML classification methods. The aim here is not to compare ML models per se but rather to demonstrate that the model parameters alone provide sufficient information for accurate decoding, regardless of the chosen ML model.

To ensure consistency, only neurons that responded to both stimulus intensities were considered, allowing for a robust train and test set derived from the same neuronal populations. Performance metrics for each model after hyperparameter tuning through grid search are reported in Tables 8 and 9. These tables summarize the models’ F1, accuracy, precision, recall, and ROC AUC scores, computed using macro averaging to handle potential class imbalances. For MO cortex stimulation, the test set included 14 neurons per class, while for SS cortex stimulation, it included 17 neurons per class.

**Table 8:**
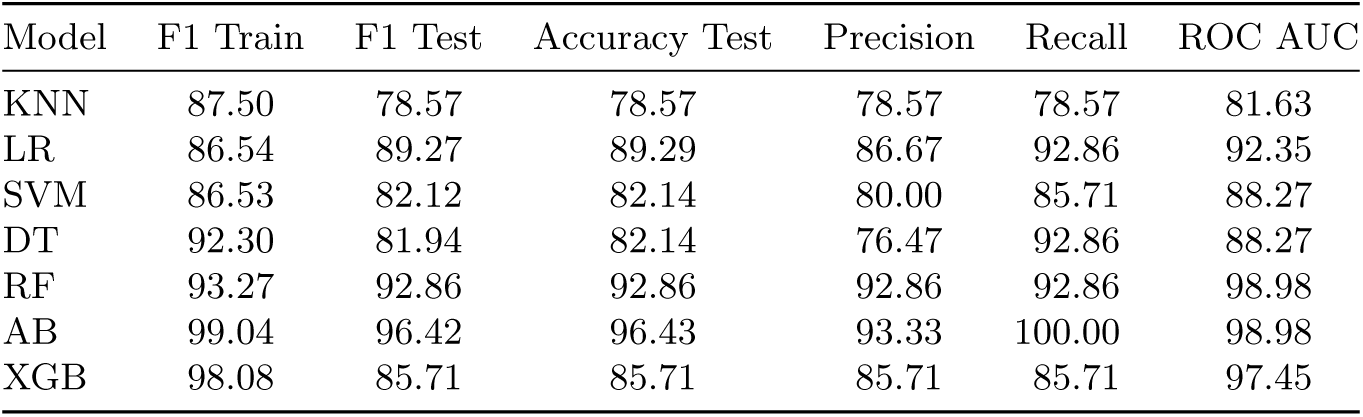
Performance metrics for ML models—K-nearest neighbors (KNN), Logistic Regression (LR), Support Vector Machines (SVM), Decision Tree (DT), Random Forest (RF), AdaBoost (AB), and XGBoost (XGB): stimulus intensity decoding in MO cortex stimulation.

| Model | F1 Train | F1 Test | Accuracy Test | Precision | Recall | ROC AUC |
| --- | --- | --- | --- | --- | --- | --- |
| KNN | 87.50 | 78.57 | 78.57 | 78.57 | 78.57 | 81.63 |
| LR | 86.54 | 89.27 | 89.29 | 86.67 | 92.86 | 92.35 |
| SVM | 86.53 | 82.12 | 82.14 | 80.00 | 85.71 | 88.27 |
| DT | 92.30 | 81.94 | 82.14 | 76.47 | 92.86 | 88.27 |
| RF | 93.27 | 92.86 | 92.86 | 92.86 | 92.86 | 98.98 |
| AB | 99.04 | 96.42 | 96.43 | 93.33 | 100.00 | 98.98 |
| XGB | 98.08 | 85.71 | 85.71 | 85.71 | 85.71 | 97.45 |

**Table 9:** Performance metrics for ML models: stimulus intensity decoding in SS cortex stimulation.

| Model | F1 Train | F1 Test | Accuracy Test | Precision | Recall | ROC AUC |
| --- | --- | --- | --- | --- | --- | --- |
| KNN | 68.68 | 67.62 | 67.65 | 66.67 | 70.59 | 69.20 |
| LR | 68.74 | 64.58 | 64.71 | 66.67 | 58.82 | 67.47 |
| SVM | 76.56 | 67.39 | 67.65 | 65.00 | 76.47 | 76.82 |
| DT | 88.28 | 61.46 | 61.76 | 64.29 | 52.94 | 65.57 |
| RF | 91.41 | 70.49 | 70.59 | 68.42 | 76.47 | 71.63 |
| AB | 100.00 | 67.62 | 67.65 | 66.67 | 70.59 | 67.13 |
| XGB | 100.00 | 64.21 | 64.71 | 61.90 | 76.47 | 66.44 |

Overall, the results highlight that the parameters of the underlying model effectively support stimulus intensity decoding. Among the models, Random Forest (RF) showed the highest performance and was thus selected for assessing feature importance. The analysis, based on the Mean Decrease in Impurity method (MDI), revealed that for the decoding task, the parameters associated with the second Gaussian component (particularly its mean and standard deviation) were the most influential features. This trend is especially pronounced in the case of SS cortex stimulation, as depicted in Figures 17 and 18.

**Figure 17:**
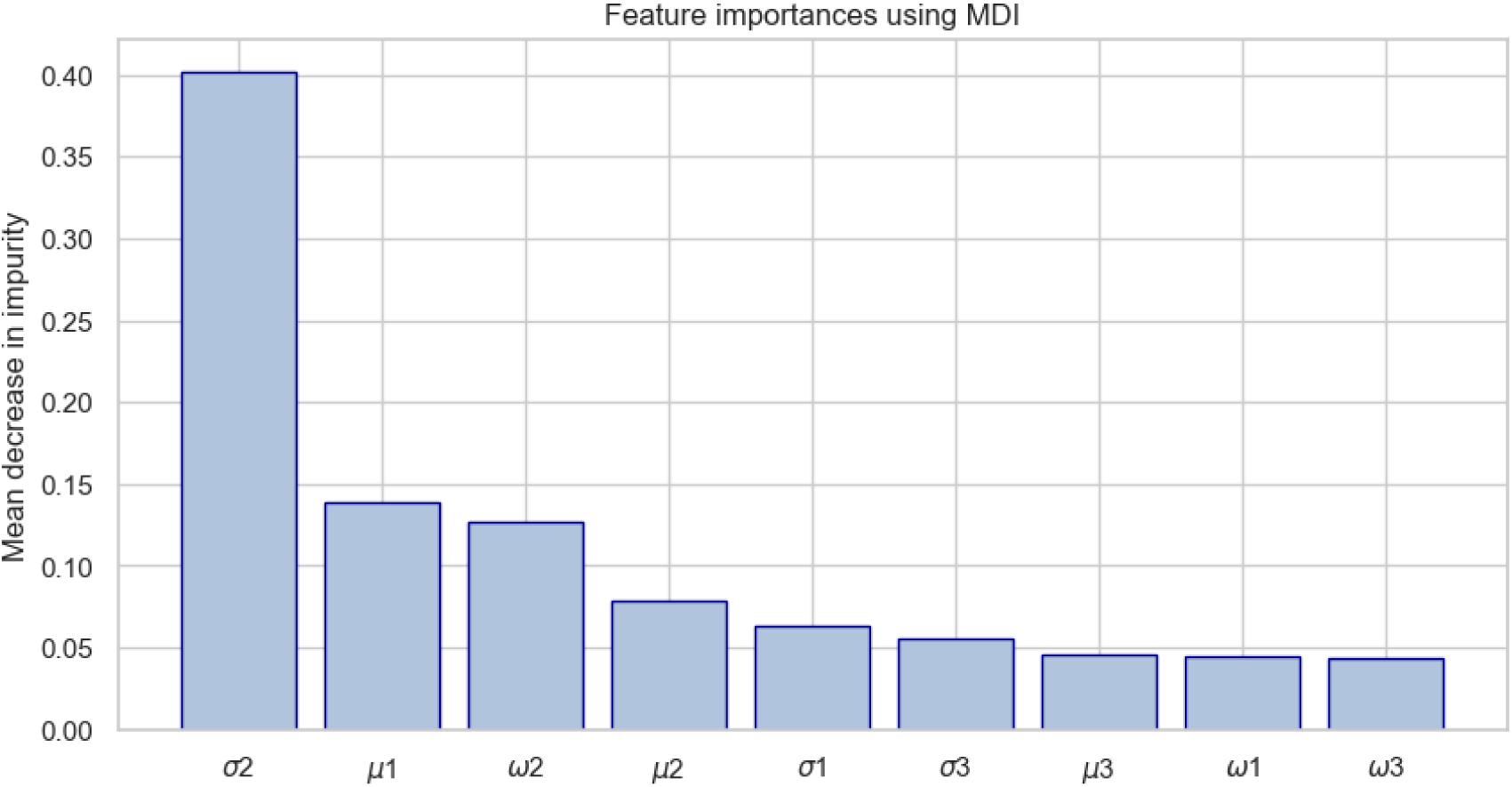
Feature importance of RF Model for stimulus intensity decoding in MO cortex. The feature with the highest importance is the standard deviation of the first curve describing the late response (i.e. *σ*_2_).

**Figure 18:**
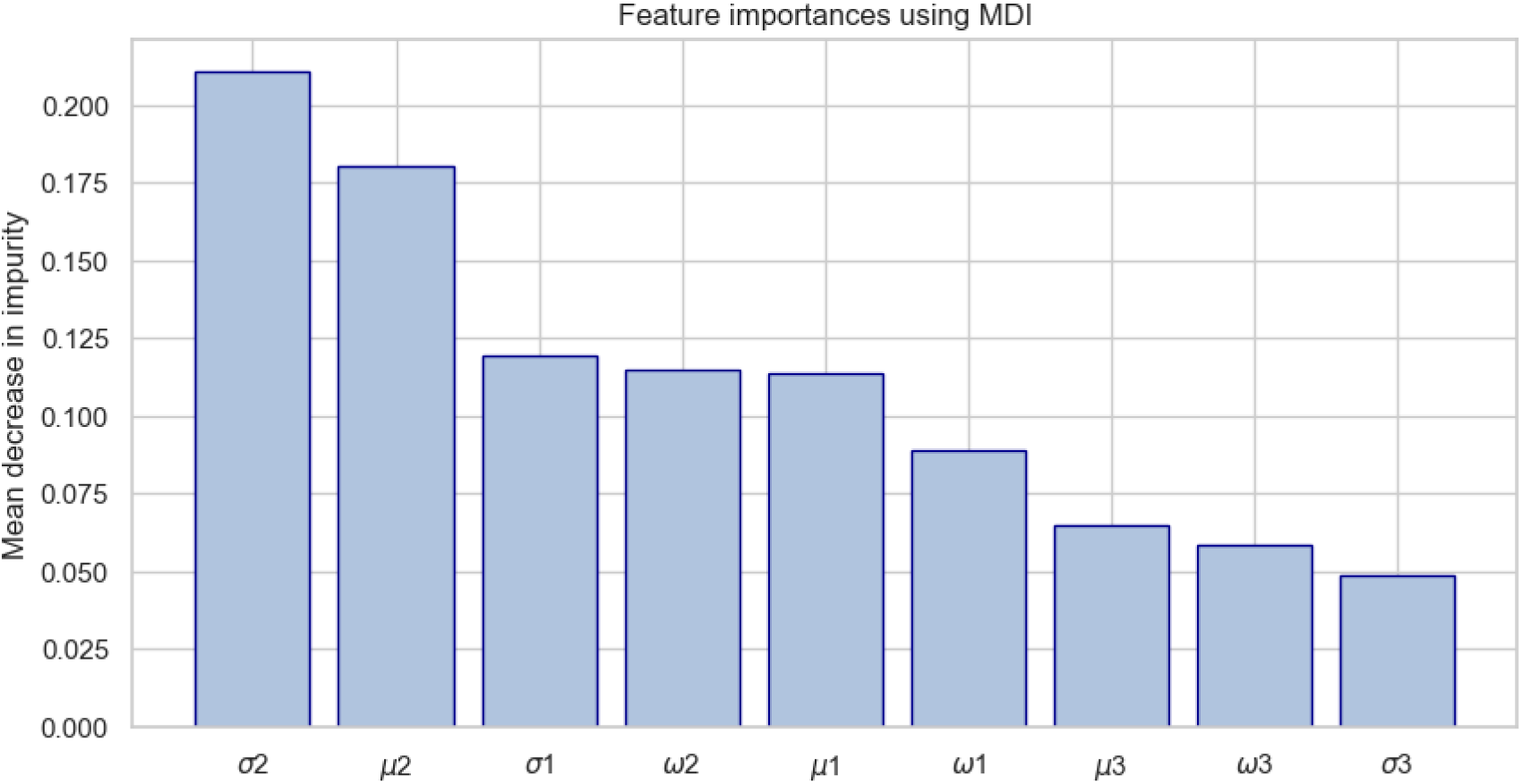
Feature importance of RF model for stimulus intensity decoding in SS cortex. The features with the highest importance are the standard deviation and the mean of the first curve describing the late response (i.e. *σ*_2_ and *µ*_2_).

These findings suggest that the second Gaussian component parameters, particularly the standard deviation, play a pivotal role in identifying stimulus intensity, highlighting the effectiveness of this model’s parameters for decoding. This insight underscores the robustness of the model’s parameterization for capturing relevant neural response features that facilitate accurate decoding of stimulus characteristics.

### 3.5 Brain Area Decoding

We examined the classification of neural responses across three brain areas—SS cortex, MO cortex, and SMTH—following electrical stimulation applied either to the MO cortex or the SS cortex. This analysis was conducted using datasets corresponding to the highest intensity stimulus, which allows for a clearer differentiation of neural activity across the regions.

For MO cortex stimulation, test support consisted of 84 neurons from SMTH, 62 from MO cortex, and 16 from SS cortex; for SS cortex stimulation, test support was 21 from SMTH, 50 from MO cortex, and 32 from SS cortex. This distribution impacts the evaluation, particularly in classes with fewer samples, influencing the stability of precision, recall, and F1 score. The performance metrics, shown in Tables 10 and 11, include F1 macro score, accuracy, precision, recall, and ROC AUC, all calculated with macro averaging to address class imbalance and give equal importance to each class.

**Table 10:**
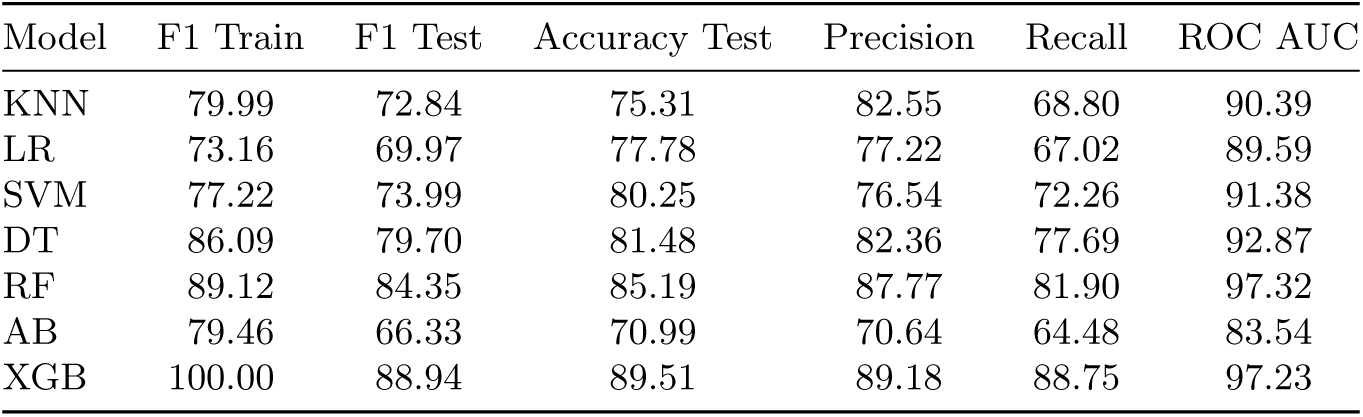
Performance metrics for ML models - KNN, LR, SVM, DT, RF, AB, and XGB: classification of brain areas when the MO cortex is stimulated. The metrics are computed using macro averaging.

**Table 11:** Performance metrics of ML models - classification of brain areas when SS cortex is stimulated. The metrics are computed with macro averaging.

| Model | F1 Train | F1 Test | Accuracy Test | Precision | Recall | ROC AUC |
| --- | --- | --- | --- | --- | --- | --- |
| KNN | 78.37 | 68.85 | 72.82 | 70.61 | 67.83 | 84.83 |
| LR | 48.13 | 51.14 | 60.19 | 55.21 | 52.35 | 78.48 |
| SVM | 50.10 | 47.87 | 65.05 | 44.49 | 52.92 | 85.79 |
| DT | 85.57 | 71.84 | 73.79 | 71.56 | 72.21 | 85.00 |
| RF | 88.43 | 79.27 | 82.52 | 85.70 | 76.54 | 91.66 |
| AB | 67.62 | 61.53 | 61.17 | 63.16 | 62.22 | 73.93 |
| XGB | 100.00 | 76.71 | 78.64 | 77.45 | 76.09 | 89.86 |

The primary result here is not a comparison among ML algorithms, but rather the demonstration that the model’s parameters alone, regardless of the chosen ML algorithm, are sufficient for decoding neural responses with high accuracy. The success of these parameters in achieving strong classification performance confirms that they encapsulate the essential characteristics needed to distinguish neural activity across brain areas. The RF and XGB models achieved particularly high performance, with RF showing an F1 test score of 84.35 and ROC AUC of 97.32 under MO cortex stimulation, and similar strong results under SS cortex stimulation.

Feature importance analysis for the RF model, illustrated in Figures 19 and 20, further highlights that among all parameters, those describing the first Gaussian component (mean and standard deviation) are the most predictive for this task. This finding underscores the central role of these initial Gaussian parameters in capturing key neural response patterns for effective brain area classification.

**Figure 19:**
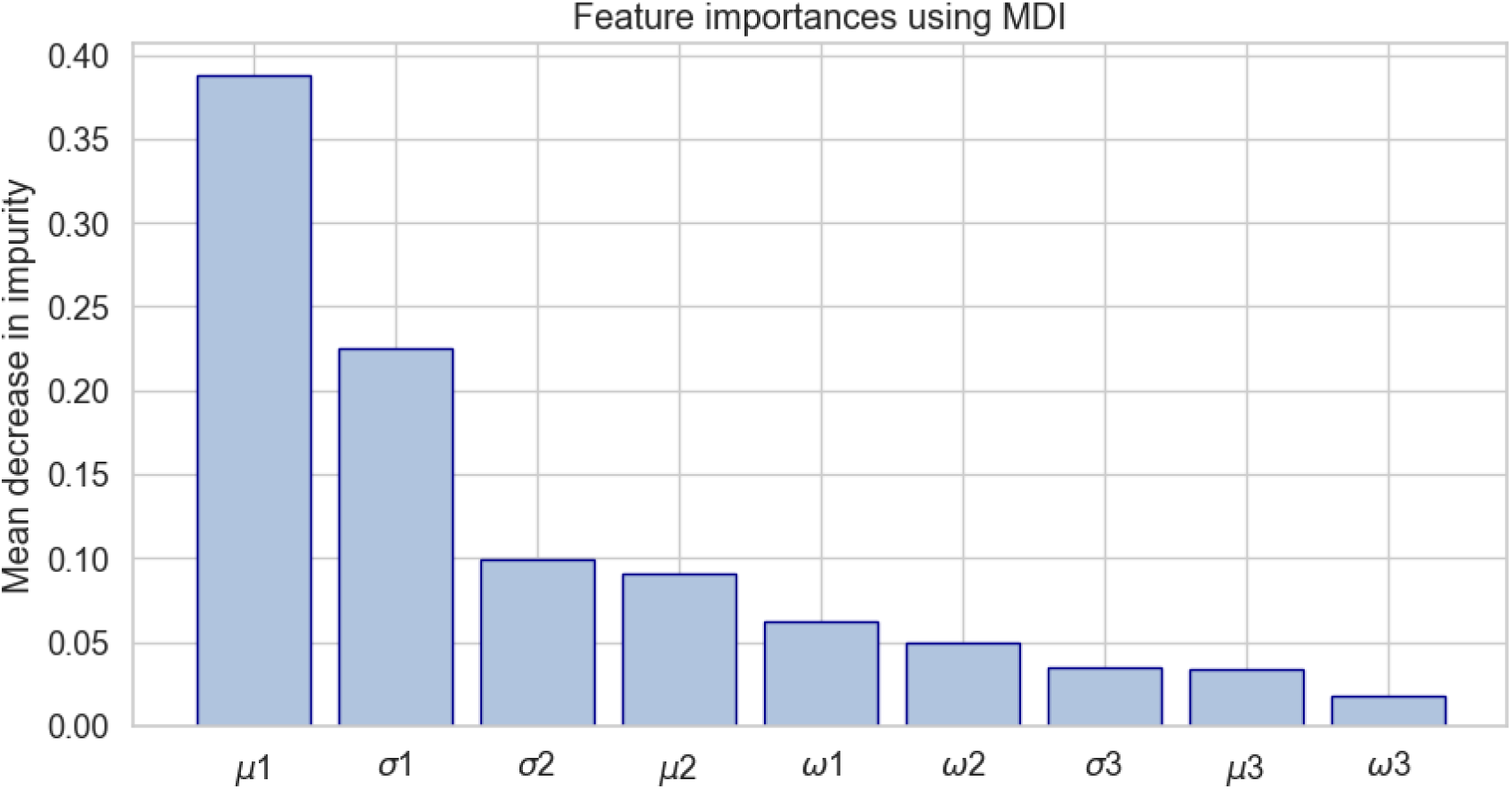
Feature importance of the RF model for classification of brain areas when the MO cortex is stimulated. The feature with the highest importance is the mean of the curve describing the early response (i.e. *µ*_1_).

**Figure 20:**
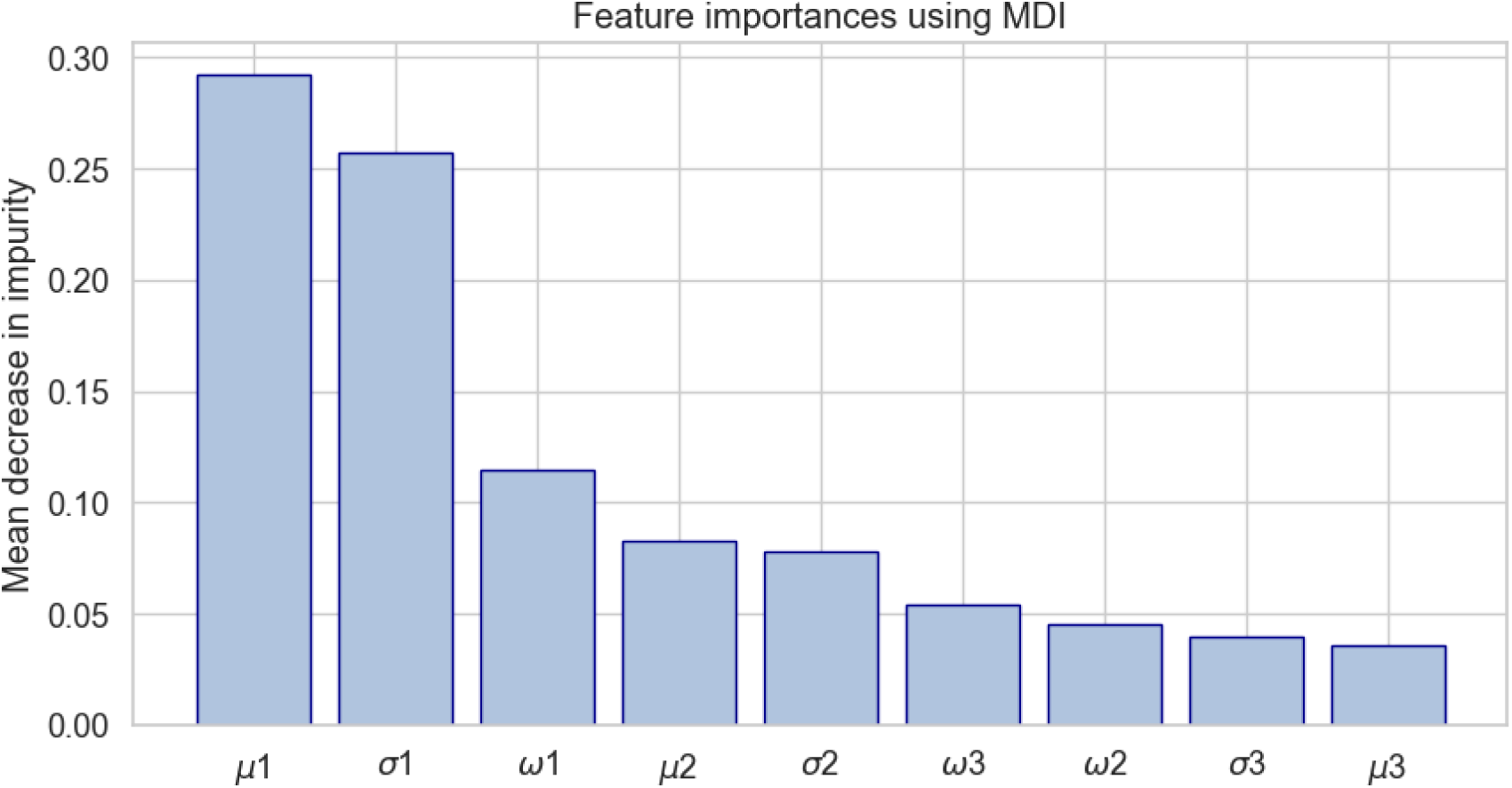
Feature importance of the RF model for classification of brain areas When the SS cortex is stimulated. The features with the highest importance are the mean and the standard deviation of the curve describing the early response (i.e. *µ*_1_ and *σ*_1_).

## 4 Discussion

### 4.1 Statistical Modeling of Electrically-Evoked Neural Response

#### 4.1.1 Modeling Neural Responses to Electrical Stimulation

The purpose of this study is to implement a statistical framework for characterizing the neural responses evoked in mice cortex and thalamus from the electrical stimulation of the cortex. Our framework combines Bayesian theory with Dirichlet mixture models to model in a time-resolved fashion the neuronal spiking evoked by electrical cortical stimulation, as described by the Peri-Stimulus Time Histogram (PSTH). For this purpose, we analyzed the dataset published by Claar et al. [22] that consists of invasive electrophysiological recordings performed through Neuropixel probes from motor (MO) and somatosensory (SS) cortex and sensorimotor related thalamic nuclei (SM-TH) during the electrical stimulation of the MO and SS cortex. In line with previous studies, we found that the response evoked within the stimulated cortex (either MO or SS cortex) and SM-TH reveals a triphasic pattern that can be divided into an early (0-50ms) and a late response (50-300ms), separated by an off period. The early response consists of a brief spiking induced by the local stimulation, while the late response reflects a rebound excitation [22]. We applied our framework to model and capture the temporal dynamics of the neuronal responses evoked in these three brain areas by MOs and SS stimulation. We applied several models, including Gaussian and Inverse Gaussian mixture probability distributions, and tested their goodness of fit using the Kolmogorov-Smirnov (KS) test. The model achieving the best performance was a Dirichlet Mixture Model consisting of three Gaussian distributions (one for the early response and the other two for the late response) described by the mean (*µ_i_*), the standard deviation (*σ_i_*), and the weight (*ω_i_*) associated with each (the *i*-th) Gaussian curve (see Table 7). It is worth mentioning that the model’s performance depends on the repeated presentation of the same stimulus, which may limit its applicability in contexts where stimuli are varied or unpredictable

#### 4.1.2 Characterization and Comparison of Response Features

We assessed how the model parameters can distinguish different stimulus intensities (i.e. low and high) and neurons belonging to different brain areas (i.e. MO and SS cortex and SM-TH) (see section 2.5). We applied the model exclusively to responsive neurons (see Section 2.2). However, it should be noted that the proposed method identifies neurons responsive to electrical stimuli without differentiating between early (0–50 ms) and late (50–300 ms) responses. Therefore, neurons active in either or both time windows were included in the analysis.

First, we tested whether different electrical stimulation intensities and brain areas were associated with different model parameters. We found significant differences between the response features evoked by electrical stimulation with high and low intensity in 3 out of 6 features for MO cortex stimulation (Figure 8 and Table 12) and in 2 out of 6 features for SS cortex stimulation (Figure 10 and Table 13). Additionally, we identified significant differences in the pairwise comparison of the model features across the three brain areas. In the response evoked by MO stimulation, from 3 to 4 features over 6 were significantly different when comparing each pair of regions (Figure 12 and Table 14). In contrast, the response to SS stimulation shows fewer significant differences (e.g. in the comparison between MO cortex and SMTH, no features are significantly different) (Figure 13 and Table 15). This may be attributed to lower statistical power resulting from the smaller sample size.

We further leveraged the model by using machine learning algorithms to classify both the stimulation intensity and the specific brain area of each neuron. The high performance achieved—an F1 score of 92.86% for stimulus intensity decoding and 84.35% for brain region classification in datasets where the MO cortex is stimulated—demonstrates the model’s ability to effectively distinguish between different stimulus intensities and brain areas in this condition. Performance remains high but is lower when the SS cortex is stimulated, with F1 scores of 70.49% for stimulus intensity decoding and 79.27% for brain region classification. Therefore, given the statistical tests and machine learning results, the proposed framework accurately distinguishes all classes when the MO cortex is stimulated, whereas classification performance worsens when stimulating the SS cortex. This is likely due to differences in the thickness of the cortical layers in these areas. Specifically, stimulation of the MO reaches layer 5, while stimulation of the SS reaches layer 4. This occurs because layer 4 in the SS is thicker than in the MO, meaning that even when stimulating at the same depth, we activate different layers: layer 4 in the SS and layer 5 in the MO. Notably, layer 4 primarily supports intra-cortical projections, whereas stimulation of layer 5 results in massive projections to the thalamus [32, 33].

#### 4.1.3 Biological Validation and Alignment with Previous Findings

To ensure that our model’s features captured biologically relevant properties, we verified that they aligned with previous findings from Claar et al. [22], reporting that the median first spike latency following stimulus onset was shorter in the stimulated cortex than in the thalamus during the early response, with the trend reversing in the late response. We computed the median values of *µ*_1_ and *µ*_2_, corresponding to the mean of the early and first part of the late response, respectively, for each subject. As shown in Figure 16, our findings mirror those of Claar et al., so that in the early response, the stimulated cortex exhibits the mean latency (*µ*_1_) before SM-TH, while in the late response, the stimulated cortex shows mean latency (*µ*_2_) larger than SM-TH. This correspondence indicates how our model extracts biologically relevant features that align with the existing knowledge about the underlying physiology of the system.

### 4.2 Neural Characterization: Insights

By analyzing the distribution of the model features, we identified consistent patterns and relationships in neural activity, which provided interesting neurobiological insights.

#### 4.2.1 Neural Response features distinguish across stimulation intensities

The threshold method used to count the number of bins with marked responses shows that a smaller percentage of neurons exhibit a pronounced early response to low stimulation compared to high stimulation in the MO cortex. Conversely, the SS cortex shows no significant difference in early response between low and high stimulation intensities. This difference between SS and MO stimulation may reflect several mechanisms, such as a saturation of the recorded neurons in SS. On the other hand, the MO cortex results suggest that low-intensity stimulation may not be sufficient to activate all recorded neurons in this region, as not all responding neurons show a pronounced early response as they do at high intensity. We know that cortical neurons that are activated from the stimulus engage the thalamus, which in turn engages in bursting activity that will trigger again cortical neurons (the late response is influenced by the thalamus activity [22, 23]). Therefore, for low stimulation intensity, neurons which were not initially activated by electrical stimulation in the early response show a response in the late stage due to thalamus-mediated inputs.

The most important model parameter for intensity decoding is the standard deviation of the first part of the late response (i.e. *σ*_2_). One potential explanation is that for low stimulation intensities, the stimulus poorly engages the stimulated cortex, which will not provide enough inputs to induce strong thalamic bursts, resulting in more variability in the latency of the rebound excitation.

These insights suggest that when the stimulus intensity is low, the reduced activation of the neural pathways may result in less reliable and more variable signal transmission, thus making the late response less predictable. This increased variability at low intensities can impact the overall accuracy and reliability of the neural response, highlighting the importance of sufficient stimulus strength for eliciting consistent neural responses.

#### 4.2.2 Neural response features distinguish stimulated and non-stimulated cortical areas

We observed similar properties in the early response of the stimulated area, independently of whether the stimulation is administered to SS and MO (Figures 14, 15). Indeed, the early responses of the stimulated site, either SS or MO, constantly show brief latency (i.e. small values of *µ*_1_) and a low variability (i.e. small values of *σ*_1_) compared to the non-stimulated site. While more variable, the early response latency and variability of the non-stimulated site are correlated. Therefore, neurons whose early response shows higher mean latency will tend to show also more variability. A potential explanation for our finding is that neurons in the non-stimulated cortex receive the stimulus transynaptically (either through one or more synapses) from the stimulated cortex, and that these synaptic stations introduce a delay and variability in the neural response evoked in the non-stimulated cortex. This correlation was absent in the late response, only, *µ*_2_ presents higher values for the stimulated cortex and lower for the non-stimulated one.

Regarding the brain area classification between MO and SS cortex and SMTH, the features with highest importance are the mean latency and standard deviation of the early response (i.e. *µ*_1_ and *σ*_1_) for both datasets. Indeed, the stimulated cortex exhibits a sharper early response compared to the non-stimulated one, while the SMTH early response occurs at higher latencies with respect to the stimulated cortex.

## 5 Conclusion

This work presents a statistical framework for characterizing neural responses to electrical stimuli in the brain cortex and the thalamus. Bayesian Theory and Dirichlet Mixture Models were employed to model the neuronal electrically evoked response by computing the probability of spike occurrence at specific latencies from stimulus onset. The proposed model exhibits sufficient flexibility and robustness to accurately capture the observed data. It aligns with the constraints of the available data, maintaining feasibility without excessive complexity. Notably, the model remains interpretable from a physiological perspective and aligns with existing knowledge about the underlying physiology of the system. In fact, the observed early and late response patterns are consistent with previous studies. This encoding approach was leveraged using machine learning methods for stimulus decoding and brain zone classification. The Random Forest classifier achieved an F1 score of 92.86% for stimulus intensity decoding and 84.35% for brain region classification.

This study highlights how our statistical model features offer key insights into neural responses to electrical stimulation, shedding light on cortical-thalamic dynamics and advancing neuroscience research and neural interface technologies. Indeed, we observed that neurons in the stimulated cortex exhibit an early, sharp response, while those in non-stimulated cortex display a more variable, delayed activation, likely due to transynaptic transmission. Moreover, our analysis revealed distinct response patterns depending on stimulus intensity: higher intensities evoked more precise early and late responses, whereas lower intensities led to greater variability.

Future work should aim to enhance model validation and explore its applicability to a wider range of neural responses and experimental conditions.

## Acknowledgments

We would like to thank Irene Rembado, Leslie D. Claar, Lydia Marks, and Christof Koch for providing the data and the insightful discussions.

## Funding

This work has been supported by:

- Italian Ministry of Health, Foundation IRCCS Ca’ Granda Ospedale Maggiore Policlinico Grant Ricerca Corrente 2024;
- PNC “Hub Life Science-Diagnostica Avanzata (HLS-DA), PNC-E3-2022-23683266– CUP: C43C22001630001”;
- European Union - Next Generation EU, Mission 4, Component 2, CUP B93D21010860004”, Spoke 3;
- The Italian Ministry of Education and Research (MUR): Dipartimenti di Eccellenza Program 2023–2027 -Dept. of Pathophysiology and Transplantation, University of Milan’.
- Manava Plus srl.

## Conflict of interest

Diana Nigrisoli and Nicolas Seseri are junior scientists of Manava Plus srl, Simone Russo is the Chief Medical Officer of Manava Plus srl, Ruggero Freddi is a senior researcher of Manava Plus srl.

## Appendix

**Figure 21:**
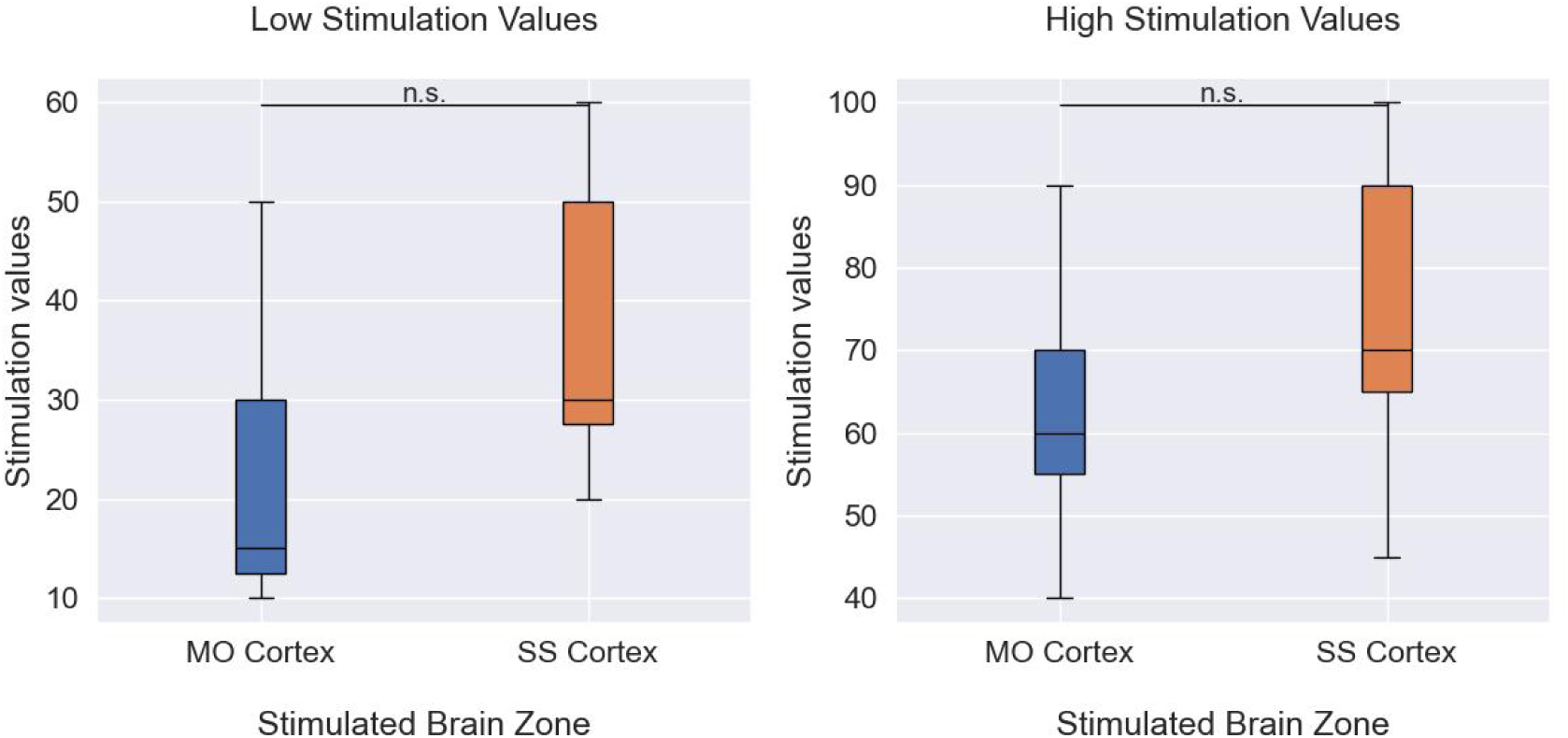
Distribution of current stimulation intensity values across all subjects of Claar et al. experiment. The median and IQR (inter-quartile range: 25th and 75th percentiles) are reported. The outcome of the statistical analysis with Mann-Whitney tests: n.s. no significant evidence to reject null hypothesis (*p >* 0.05), * weak evidence to reject null hypothesis (0.01 *< p <* 0.05), ** strong evidence to reject null hypothesis (0.001 *< p <* 0.01), and *** very strong evidence to reject null hypothesis (*p <* 0.001).

**Table 12:** Model parameters derived to describe neuronal responses in the motor (MO) cortex following electrical stimulation within the same cortical area. The median and IQR (inter-quartile range: 25th and 75th percentiles) of the features values are reported for every stimulation intensity: low stimulation intensities (LS) and high stimulation intensities (HS). The outcome of the statistical analysis with Wilcoxon signed-rank test between the classes is given by the p-value: n.s. no significant evidence to reject null hypothesis (*p >* 0.05), * weak evidence to reject null hypothesis (0.01 *< p <* 0.05), ** strong evidence to reject null hypothesis (0.001 *< p <* 0.01), and *** very strong evidence to reject null hypothesis (*p <* 0.001).

| parameter | LS | HS | LS vs HS |
| --- | --- | --- | --- |
| $\mu_1$ | 0.006 [0.004 - 0.043] | 0.005 [0.004 - 0.005] | *** |
| $\mu_2$ | 0.17 [0.154 - 0.185] | 0.174 [0.165 - 0.185] | ns |
| $\mu_3$ | 0.264 [0.247 - 0.277] | 0.251 [0.235 - 0.264] | * |
| $\sigma_1$ | 0.002 [0.001 - 0.025] | 0.001 [0.001 - 0.002] | ns |
| $\sigma_2$ | 0.049 [0.042 - 0.055] | 0.018 [0.012 - 0.029] | *** |
| $\sigma_3$ | 0.021 [0.013 - 0.027] | 0.026 [0.021 - 0.034] | ns |
| $\omega_1$ | 0.177 [0.093 - 0.26] | 0.24 [0.16 - 0.348] | - |
| $\omega_2$ | 0.58 [0.453 - 0.654] | 0.371 [0.221 - 0.505] | - |
| $\omega_3$ | 0.225 [0.13 - 0.369] | 0.308 [0.211 - 0.458] | - |

**Table 13:** Model parameters derived to describe neuronal responses in the somatosensory (SS) cortex following electrical stimulation within the same cortical area. The median and IQR (inter-quartile range: 25th and 75th percentiles) of the features values are reported for every stimulation intensity: low stimulation intensities (LS) and high stimulation intensities (HS). The outcome of the statistical analysis with Wilcoxon signed-rank test between the classes is given by the p-value: n.s. no significant evidence to reject null hypothesis (*p >* 0.05), * weak evidence to reject null hypothesis (0.01 *< p <* 0.05), ** strong evidence to reject null hypothesis (0.001 *< p <* 0.01), and *** very strong evidence to reject null hypothesis (*p <* 0.001).

| parameter | LS | HS | LS vs HS |
| --- | --- | --- | --- |
| $\mu_1$ | 0.005 [0.004 - 0.017] | 0.004 [0.003 - 0.005] | ns |
| $\mu_2$ | 0.172 [0.154 - 0.192] | 0.188 [0.173 - 0.203] | *** |
| $\mu_3$ | 0.266 [0.255 - 0.277] | 0.264 [0.247 - 0.279] | ns |
| $\sigma_1$ | 0.002 [0.001 - 0.01] | 0.001 [0.001 - 0.002] | ns |
| $\sigma_2$ | 0.045 [0.031 - 0.055] | 0.036 [0.018 - 0.047] | *** |
| $\sigma_3$ | 0.02 [0.013 - 0.027] | 0.021 [0.013 - 0.031] | ns |
| $\omega_1$ | 0.175 [0.105 - 0.252] | 0.178 [0.085 - 0.349] | - |
| $\omega_2$ | 0.615 [0.429 - 0.706] | 0.476 [0.221 - 0.653] | - |
| $\omega_3$ | 0.212 [0.137 - 0.321] | 0.22 [0.109 - 0.42] | - |

**Table 14:** Model parameters derived to describe neuronal responses in SM-TH, MO cortex, and SS cortex neurons when MO cortex is electrically stimulated with high current intensities. The median and IQR (inter-quartile range: 25th and 75th percentiles) of the features values are reported for every brain area. The outcome of the statistical analysis with Mann-Whitney tests: n.s. no significant evidence to reject null hypothesis (*p >* 0.05), * weak evidence to reject null hypothesis (0.01 *< p <* 0.05), ** strong evidence to reject null hypothesis (0.001 *< p <* 0.01), and *** very strong evidence to reject null hypothesis (*p <* 0.001).

| parameter | MO cortex | SS cortex | SM-TH | MO vs SS | MO vs SM-TH | SS vs SM-TH |
| --- | --- | --- | --- | --- | --- | --- |
| $\mu_1$ | 0.005 [0.004 - 0.006] | 0.028 [0.019 - 0.048] | 0.009 [0.008 - 0.01] | *** | *** | *** |
| $\mu_2$ | 0.172 [0.163 - 0.186] | 0.151 [0.128 - 0.177] | 0.16 [0.149 - 0.178] | *** | *** | ns |
| $\mu_3$ | 0.25 [0.237 - 0.265] | 0.263 [0.244 - 0.273] | 0.25 [0.233 - 0.267] | ns | ns | ns |
| $\sigma_1$ | 0.001 [0.001 - 0.002] | 0.014 [0.008 - 0.024] | 0.003 [0.002 - 0.004] | *** | *** | *** |
| $\sigma_2$ | 0.017 [0.011 - 0.026] | 0.047 [0.036 - 0.053] | 0.024 [0.018 - 0.036] | *** | *** | *** |
| $\sigma_3$ | 0.026 [0.02 - 0.033] | 0.019 [0.014 - 0.031] | 0.026 [0.02 - 0.034] | ns | ns | ns |
| $\omega_1$ | 0.233 [0.109 - 0.406] | 0.137 [0.076 - 0.186] | 0.151 [0.086 - 0.236] | - | - | - |
| $\omega_2$ | 0.363 [0.243 - 0.509] | 0.554 [0.455 - 0.674] | 0.486 [0.332 - 0.627] | - | - | - |
| $\omega_3$ | 0.323 [0.192 - 0.467] | 0.264 [0.174 - 0.395] | 0.322 [0.204 - 0.464] | - | - | - |

**Table 15:** Model parameters derived to describe neuronal responses in SM-TH, MO cortex, and SS cortex neurons When SS cortex is electrically stimulated with high current intensities. The median and IQR (inter-quartile range: 25th and 75th percentiles) of the features values are reported for every brain area. The outcome of the statistical analysis with Mann-Whitney tests: n.s. no significant evidence to reject null hypothesis (*p >* 0.05), * weak evidence to reject null hypothesis (0.01 *< p <* 0.05), ** strong evidence to reject null hypothesis (0.001 *< p <* 0.01), and *** very strong evidence to reject null hypothesis (*p <* 0.001).

| parameter | MO cortex | SS cortex | SM-TH | MO vs SS | MO vs SM-TH | SS vs SM-TH |
| --- | --- | --- | --- | --- | --- | --- |
| $\mu_1$ | 0.02 [0.012 - 0.03] | 0.004 [0.004 - 0.008] | 0.025 [0.014 - 0.041] | *** | ns | *** |
| $\mu_2$ | 0.168 [0.142 - 0.192] | 0.185 [0.169 - 0.2] | 0.153 [0.136 - 0.17] | ** | ns | *** |
| $\mu_3$ | 0.259 [0.238 - 0.271] | 0.263 [0.241 - 0.275] | 0.257 [0.247 - 0.272] | ns | ns | ns |
| $\sigma_1$ | 0.011 [0.005 - 0.019] | 0.001 [0.001 - 0.005] | 0.01 [0.005 - 0.019] | *** | ns | *** |
| $\sigma_2$ | 0.045 [0.023 - 0.054] | 0.035 [0.018 - 0.047] | 0.042 [0.029 - 0.053] | * | ns | * |
| $\sigma_3$ | 0.023 [0.015 - 0.033] | 0.022 [0.014 - 0.032] | 0.024 [0.017 - 0.031] | ns | ns | ns |
| $\omega_1$ | 0.108 [0.067 - 0.16] | 0.154 [0.063 - 0.306] | 0.116 [0.063 - 0.225] | - | - | - |
| $\omega_2$ | 0.539 [0.404 - 0.657] | 0.453 [0.237 - 0.614] | 0.543 [0.391 - 0.652] | - | - | - |
| $\omega_3$ | 0.316 [0.183 - 0.472] | 0.293 [0.123 - 0.518] | 0.291 [0.207 - 0.428] | - | - | - |

## References

[1] Jerry J Shih, Dean J Krusienski, and Jonathan R Wolpaw. “Brain-computer interfaces in medicine”. In: Mayo clinic proceedings. Vol. 87. 3. Elsevier. 2012, pp. 268–279.

[2] Sydney S Cash and Leigh R Hochberg. “The emergence of single neurons in clinical neurology”. In: Neuron 86.1 (2015), pp. 79–91.

[3] L. Paninski, J. Pillow, and J. Lewi. “Statistical models for neural encoding, decoding, and optimal stimulus design”. In: Progress in Brain Research 165 (2007), pp. 493–507.

[4] Eric R Kandel et al. Principles of neural science. Vol. 4. McGraw-hill New York, 2000.

[5] Fabrizio Gabbiani and Christof Koch. “Principles of spike train analysis”. In: Methods in neuronal modeling 12.4 (1998), pp. 313–360.

[6] Robert E Kass, Valérie Ventura, and Emery N Brown. “Statistical issues in the analysis of neuronal data”. In: Journal of neurophysiology 94.1 (2005), pp. 8–25.

[7] Wilson Truccolo et al. “A point process framework for relating neural spiking activity to spiking history, neural ensemble, and extrinsic covariate effects”. In: Journal of neurophysiology 93.2 (2005), pp. 1074–1089.

[8] R. Barbieri et al. “Construction and analysis of non-Poisson stimulus-response models of neural spiking activity”. In: Journal of Neuroscience Methods 105.1 (2001), pp. 25–37.

[9] R. Barbieri, et al. “Dynamic analyses of information encoding in neural ensembles”. In: Neural Computation 16.2 (2004), pp. 277–307.

[10] R. Levi, et al. “Bayesian supervised machine learning classification of neural networks with pathological perturbations”. In: Biomedical Physics & Engineering Express 7.6 (2021), p. 065021.

[11] Don H Johnson. “Point process models of single-neuron discharges”. In: Journal of computational neuroscience 3 (1996), pp. 275–299.

[12] Don H Johnson. “The relationship of post-stimulus time and interval histograms to the timming characteristics of spike trains”. In: Biophysical Journal 22.3 (1978), pp. 413–430.

[13] Annette M Taberner and M Charles Liberman. “Response properties of single auditory nerve fibers in the mouse”. In: Journal of neurophysiology 93.1 (2005), pp. 557–569.

[14] Jeffery T Lichtenhan and Mark E Chertoff. “Temporary hearing loss influences post-stimulus time histogram and single neuron action potential estimates from human compound action potentials”. In: The Journal of the Acoustical Society of America 123.4 (2008), pp. 2200–2212.

[15] Marko Takanen, Ian C Bruce, and Bernhard U Seeber. “Phenomenological modelling of electrically stimulated auditory nerve fibers: A review”. In: Network: Computation in Neural Systems 27.2-3 (2016), pp. 157–185.

[16] Ishita Basu, et al. “Stochastic modeling of the neuronal activity in the subthalamic nucleus and model parameter identification from Parkinson patient data”. In: Biological cybernetics 103 (2010), pp. 273–283.

[17] Roxana A Stefanescu, RG Shivakeshavan, and Sachin S Talathi. “Computational models of epilepsy”. In: Seizure 21.10 (2012), pp. 748–759.

[18] Corey J Keller, et al. “Mapping human brain networks with cortico-cortical evoked potentials”. In: Philosophical Transactions of the Royal Society B: Biological Sciences 369.1653 (2014), p. 20130528.

[19] Antonio Valentin, et al. “Single-pulse electrical stimulation identifies epileptogenic frontal cortex in the human brain”. In: Neurology 65.3 (2005), pp. 426–435.

[20] Leo Pio-Lopez, Romanos Poulkouras, and Damien Depannemaecker. “Visual cortical prosthesis: an electrical perspective”. In: Journal of Medical Engineering & Technology 45.5 (2021), pp. 394–407.

[21] L. D. Claar, et al. Simultaneous electroencephalography, extracellular electrophysiology, and cortical electrical stimulation in head-fixed mice. Available at: 10.48324/dandi.000458/0.230317.0039. 2023.

[22] L. D. Claar, et al. “Corticothalamo-cortical interactions modulate electrically evoked EEG responses in mice”. In: Elife 12 (2023), RP84630.

[23] S. Russo, et al. “Thalamic feedback shapes brain responses evoked by cortical stimulation in mice and humans”. In: bioRxiv (2024), pp. 2024–01.

[24] D. H. Johnson. “The relationship of post-stimulus time and interval histograms to the timing characteristics of spike trains”. In: Biophysical Journal 22.3 (1978), pp. 413–430.

[25] Allen Brain Atlas. Available at: https://mouse.brain-map.org/static/atlas.

[26] James J Jun, et al. “Fully integrated silicon probes for high-density recording of neural activity”. In: Nature 551.7679 (2017), pp. 232–236.

[27] M. Pachitariu, et al. “Kilosort: Real-time spike-sorting for extracellular electrophysiology with hundreds of channels”. In: BioRxiv (2016), p. 061481.

[28] Peter Barthó, et al. “Characterization of neocortical principal cells and interneurons by network interactions and extracellular features”. In: Journal of neurophysiology 92.1 (2004), pp. 600–608.

[29] Dante S Bortone, Shawn R Olsen, and Massimo Scanziani. “Translaminar inhibitory cells recruited by layer 6 corticothalamic neurons suppress visual cortex”. In: Neuron 82.2 (2014), pp. 474–485.

[30] M. D. Escobar and M. West. “Bayesian density estimation and inference using mixtures”. In: Journal of the American Statistical Association 90.430 (1995), pp. 577–588.

[31] Vance W Berger and YanYan Zhou. “Kolmogorov–smirnov test: Overview”. In: Wiley statsref: Statistics reference online (2014).

[32] Susann Boretius, et al. “MRI of cellular layers in mouse brain in vivo”. In: Neuroimage 47.4 (2009), pp. 1252–1260.

[33] Alexis M Hattox and Sacha B Nelson. “Layer V neurons in mouse cortex projecting to different targets have distinct physiological properties”. In: Journal of neurophysiology 98.6 (2007), pp. 3330–3340.

